# Multilineage plasticity in prostate cancer through expansion of stem–like luminal epithelial cells with elevated inflammatory signaling

**DOI:** 10.1101/2021.11.01.466599

**Authors:** Samir Zaidi, Jimmy L. Zhao, Joseph M. Chan, Martine P. Roudier, Kristine M. Wadosky, Anuradha Gopalan, Wouter R. Karthaus, Jungmin Choi, Kayla Lawrence, Ojasvi Chaudhary, Tianhao Xu, Ignas Masilionis, Linas Mazutis, Ronan Chaligné, Irina Linkov, Afsar Barlas, Achim Jungbluth, Natasha Rekhtman, Joachim Silber, Katia Manova–Todorova, Philip A. Watson, Lawrence D. True, Peter S. Nelson, Howard I. Scher, Dana E. Rathkopf, Michael J. Morris, Michael C. Haffner, David W. Goodrich, Dana Pe’er, Charles L. Sawyers

## Abstract

Lineage plasticity is a well–established mechanism of resistance to targeted therapies in lung and prostate cancer, where tumors transition from adenocarcinoma to small–cell or neuroendocrine carcinoma. Through single–cell analysis of a cohort of heavily–treated castration–resistant human prostate cancers (CRPC), we report a greater degree of plasticity than previously appreciated, with multiple distinct neuroendocrine (NEPC), mesenchymal (EMT–like), and other subpopulations detected within single biopsies. To explore the steps leading to this plasticity, we turned to two genetically engineered mouse models of prostate cancer that recapitulate progression from adenocarcinoma to neuroendocrine disease. Time course studies reveal expansion of stem–like luminal epithelial cells (*Sca1*+, *Psca*+, called L2) that, based on trajectories, gave rise to at least 4 distinct subpopulations, NEPC (*Ascl1*+), POU2F3 (*Pou2f3*+), TFF3 (*Tff3*+) and EMT–like (*Vim*+, *Ncam1*+)––these populations are also seen in human prostate and small cell lung cancers. Transformed L2–like cells express stem–like and gastrointestinal endoderm–like transcriptional programs, indicative of reemerging developmental plasticity programs, as well as elevated Jak/Stat and interferon pathway signaling. In sum, while the magnitude of multilineage heterogeneity, both within and across patients, raises considerable treatment challenges, the identification of highly plastic luminal cells as the likely source of this heterogeneity provides a target for more focused therapeutic intervention.

**One Sentence Summary:** Multilineage plasticity results from expansion of stem–like luminal cells with JAK/STAT activation, serving as a therapeutic target.

## Main Text

Acquired resistance to precision oncology therapies is often due to mutations in the drug target present in rare subclones, which eventually emerge during treatment (*1, 2*). This understanding has spurred the development of several next–generation targeted therapies that can overcome such “on target” resistance and extend survival (*3–5*). However, the expanded clinical use of these more effective inhibitors has led to the recognition that acquired resistance is often associated with a change in tumor histology, typified by the adenocarcinoma to neuroendocrine carcinoma transition seen in lung and prostate cancer (*6*). This histologic transition, often termed lineage plasticity, refers broadly to cell–state changes that allow cells to adapt to environmental stresses, such as those associated with invasion and metastasis, as well as to the selective pressure of drug therapy (*7*). Work across a range of model systems has shown that transcriptional programs associated with plasticity typically resemble those seen in stem and progenitor cells of various tissues and in various developmental pathways (*8–10*). Unlike the more conventional mechanism of drug resistance due to “on target” mutation, lineage plasticity does not appear to be associated with the acquisition of new mutations; however, mutational contexts such as loss of tumor suppressor genes can predispose to acquiring a state of plasticity (*11–14*).

To better understand the process by which plasticity begins and subsequently evolves in a common human tumor, we used single cell genomics to characterize a cohort of early and late stage human prostate cancers and two genetically engineered mouse prostate cancer models (GEMM) that undergo an adenocarcinoma to neuroendocrine lineage transition (*9*) (Fig. 1A). Integrative computational analysis of these mouse and human datasets, together with human small cell lung cancer data, reveals conservation of plasticity–derived cell types and transcriptional programs, pointing toward a specific luminal subpopulation as the source of plasticity.

**Figure 1.**
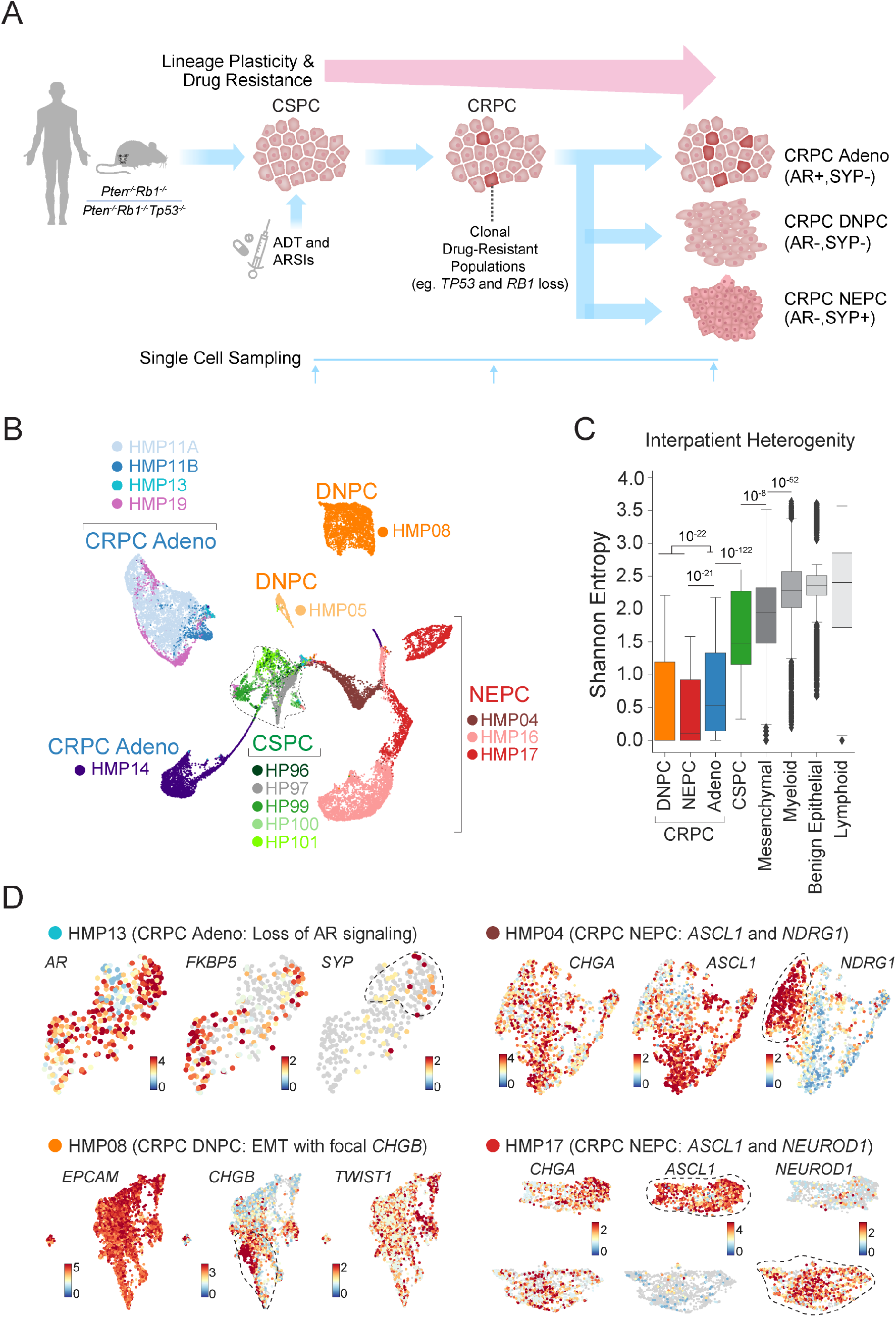
Drivers of Resistance in Metastatic Human Castration–Resistant Prostate Cancer. **(A)** Schematic of an investigative platform to explore drug resistance and lineage plasticity in early**–** and late**–**stage human prostate cancer and two autochthonous genetically engineered mouse prostate cancer models (GEMM). Human prostate cancer samples include castrate–sensitive prostate cancer (CSPC) and castrate–resistant prostate cancer (CRPC). The latter includes CRPC adenocarcinoma (AR+, SYP–), double–negative prostate cancer (DNPC) (AR–, SYP–), and transformed neuroendocrine prostate cancer (NEPC) (AR–, SYP+). GEMMs include organ–specific deletion (*Probasin*–Cre) of *Pten* and *Rb1* (PtR), or *Pten, Rb1,* and *Tp53* (PtRP) at relevant timepoints reflecting adenocarcinoma to NEPC transformation. **(B)** UMAPs of tumor cells (N=29,373 cells), colored by tumor ID and grouped by tumor type. **(C)** Inter–patient heterogeneity measured by Shannon entropy based on tumor frequencies. To control for cell sampling, 100 cells were subsampled from each Phenograph cluster (*k*=30) within epithelial, immune, and mesenchymal compartments 100 times with replacement (Bonferroni–adjusted Student’s t–test, **refer to Methods**). **(D)** UMAPs of individual CRPC biopsies: HMP13 (N=354 cells, CRPC adeno), HMP08 (N=4,279 cells, DNPC), HMP04 (N=1791 cells, NEPC), and HMP17 (N=2,845 cells, NEPC). For each biopsy, cells are colored based on normalized expression (log_2_X+1) for select genes demonstrating heterogenous and divergent intra–tumoral molecular resistance programs.

Recent single–cell analyses of normal human and mouse prostate tissue have defined a complex array of previously unappreciated cell types, particularly within the luminal epithelial compartment (*15–20*). We interrogated this complexity in the context of prostate cancer by first performing single–cell RNA sequencing (scRNA–seq) analysis of sixteen human samples (**Table S1**). Six patients had early–stage castration–sensitive prostate cancer (CSPC) (*2*) and ten patients had late–stage metastatic castration–resistant prostate cancer (CRPC) representing three clinically recognized phenotypes: CRPC adenocarcinoma (AR+, SYP–); double negative prostate cancer (DNPC) (AR–, SYP–); and neuroendocrine prostate cancer (NEPC) (AR–, SYP+). All three CRPC phenotypes were enriched for alterations in *TP53*, *PTEN* and *RB1* (*8, 11*) (**Table S2 and Fig. S1**).

Unsupervised clustering (*21*) on batch–corrected data was used for iterative labeling of coarse cell types into lineages using canonical markers (Fig. 1B**, Fig. S2**). Primary benign and CSPC tumor cells were labelled *per* Karthaus *et al* (*2*), whereas all epithelial cells derived from metastatic tumors were found to be malignant, except for a small benign hepatocyte population (**Fig. S2, Table S5** for DEGs**, refer to Methods**). To assess for inter–tumor heterogeneity, we determined the Shannon diversity of patient tumors (**refer to Methods**). As anticipated, CRPC samples were significantly more heterogeneous (lower entropy) than CSPC tumors (P=1.5×10^-205^, Fig. 1C). Furthermore, while CRPC samples showed minimal overlap across samples and did not adhere to *a priori* defined pathologic groupings (Fig. 1B), the CRPC adenocarcinoma subgroup showed less heterogeneity (higher entropy) than the DNPC or NEPC samples (P=1.6×10^-22^, Fig. 1C).

We next explored the factors that contribute to intra–tumoral heterogeneity as tumors displaying plasticity may harbor distinct tumor phenotypes at the *per*–biopsy level. Recurring transcriptomic trends emerged within multiple tumor biopsies including focal loss of AR signaling, gain of EMT signals, emergence of rare and early NEPC cells, and differing NEPC lineages (Fig. 1D**, Figs. S3–S4**). For example, HMP13 (CRPC adenocarcinoma) showed consistent AR expression throughout but had clusters with loss of expression of AR target genes, such as *FKBP5* and *NKX3.1*, and rare gain of *SYP* or *INSM1*+ cells, likely indicative of early transition to NEPC (Fig. 1D). HMP08, histologically classified as DNPC, showed evidence of EMT (*EPCAM+, TWIST1*+) and focal expression of the NEPC marker *CHGB* (Fig. 1D, **Fig. S4**). Two further samples, both classified histologically as NEPC, displayed either distinct clusters of *ASCL1*+ and *NDRG1*+ (an EMT marker) (HMP04, Fig. 1D, **Fig. S4**) or *ASCL1*+ and *NEUROD1*+ cells (HMP17, Fig. 1D) that, in human small cell lung cancer, are considered distinct disease subtypes. Collectively, these cases (each of which serves as its own case study) illustrate the complexity of aberrant lineages that can emerge from a relatively uniform population of adenocarcinoma cells in response to the selective pressure of androgen receptor signaling inhibitors (ARSIs).

To gain more insight into the early stages of lineage plasticity, we turned to two prostate GEMMs that model the adenocarcinoma to NEPC transition following prostate**–**specific deletion of the human**–**relevant tumor suppressor genes *PTEN*, *RB1* and *TP53*. scRNA–seq was performed on whole prostates harvested from 29 mice [9 wildtype (WT); 7 *Pten*^-/-^*Rb*^-/-^ (PtR); 13 *Pten*^-/-^*Rb*^-/-^*Trp53*^-/-^ (PtRP)] at relevant timepoints reflecting adenocarcinoma to NEPC transformation (Fig. 2A**, Fig. S5**). Transcriptomes from 67,622 cells were analyzed using *Gfp* to mark those that underwent Cre recombination and visualized by UMAP, with cell types identified by coarse labelling (Fig. 2B**, Fig. S5, refer to Methods**). In addition to the expected adenocarcinoma (*Epcam***–***Gfp*) and NEPC (*Syp+*) populations, we were surprised to see three additional *Gfp*+ clusters defined by expression of *Pou2f3* and *Dclk1* (*Pou2f3–Gfp*), *Vim, Twist2* and *Ncam1* (EMT*–Gfp*), and *Tff3* (*Tff3–Gfp*), respectively. POU2F3 is a transcription factor (TF) expressed by rare chemosensory cells in normal gut and lung tissue (often called tuft cells), whereas *Tff3* encodes a secreted protein expressed by intestinal columnar epithelial cells involved in maintaining mucosal integrity. Of note, POU2F3 also defines a non–neuroendocrine variant subtype of lung cancer with morphologic features of small cell carcinoma distinct from ASCL1 and NEUROD1 subtypes (*12, 22*).

**Figure 2.**
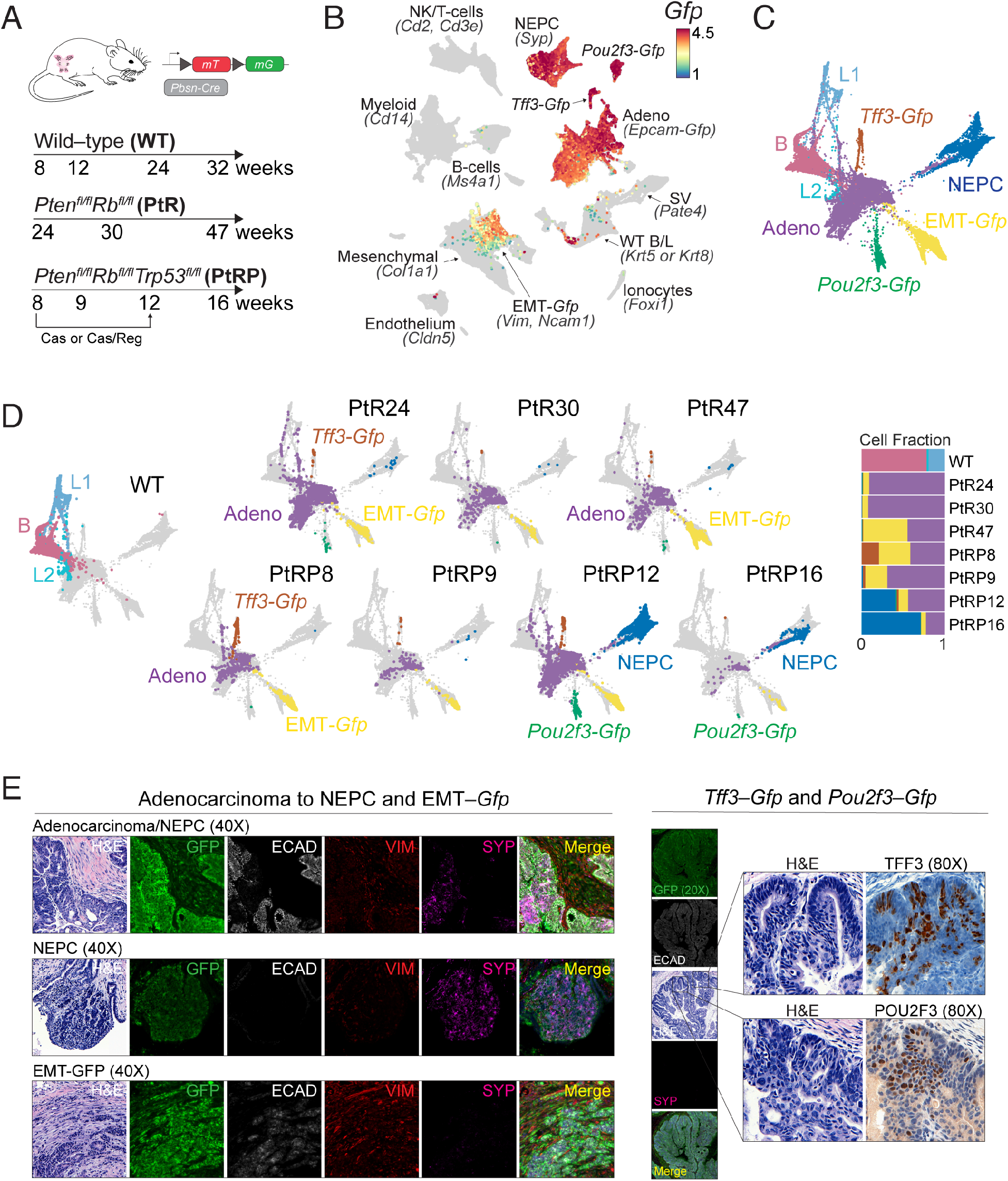
Modeling Multilineage Plasticity in GEMMs. **(A)** Experimental design includes WT (9 mice), PtR (7 mice, *PbCre*:*Rosa26*^mT/mG^*Pten*^fl/fl^*Rb1*^fl/fl^), and PtRP (13 mice, *PbCre*: *Rosa26*^mT/mG^*Pten*^fl/fl^*Rb1*^fl/fl^*Tp53*^fl/fl^) at labelled time points. **(B)** UMAP of GEMMs (N=67,622 cells) shown for all cell types based on imputed *Gfp* expression (restricted to non–immune cells) as a marker for mutant cells (**refer to Methods**). **(C)** Mutant *Gfp*–positive and wild–type cells (N=28,934) were subsetted. Shown is a force–directed layout (FDL) colored by wild–type and mutant cell types. **(D)** FDLs separated by genotypes and timepoints and colored by cell types (left, WT with N=7,435 cells; top PtRP 24/30/47 weeks with N=2,565/662/1,441 cells; bottom PtRP 8/9/12/16 weeks with N=981/353/4,984/569 cells) with stacked bar plot of fraction of cell types *per* genotype and time point (right). **(E)** Validation of mutant subtypes, including adenocarcinoma to NEPC transition, NEPC, and EMT by hematoxylin/eosin and multiplex immunofluorescence (mIF) (40x). Shown are GFP (mG, green), DAPI (nuclei, blue), VIM (mesenchymal, red), SYP (synaptophysin, purple), and E–CAD (e–cadherin, white). Also shown is mIF of adenocarcinoma state (20x) with zoomed in (80x) hematoxylin/eosin and immunohistochemistry of TFF3– and POU2F3–positive cells in mutant tissue.

We examined potential routes of plasticity by generating force**–**directed layouts (FDLs) for each genotype across different timepoints. We noted the early appearance of *Gfp*+ adenocarcinoma (*Epcam–Gfp*) as well as *Tff3–Gfp* cells, followed by the expansion of EMT–*Gfp* and NEPC and *Pou2f3–Gfp* populations, with some differences based on genotype (Figs. 2C **and** 2D). The presence of each of these subpopulations was confirmed by multiplex immunofluorescence (IF) and immunohistochemistry (IHC), which documented SYP+ (NEPC) and VIM+ (EMT) positive cells in a background of ECAD+ (or PANCK+) epithelial cells (top row), larger clusters of NEPC and EMT (middle and bottom rows) and foci of TFF3+ or POU2F3+ cells (right panel) (Fig. 2E, see **Fig. S6** for additional examples and PANCK staining).

To investigate the cellular origin of these early adenocarcinoma cells, we performed deconvolution of mixed wild**–**type epithelial phenotypes in mutant *Gfp*+ cells using Markov absorption classification (**refer to Methods**) and found the majority of cells associate with stem–like luminal 2 (*Sca1, Psca, Tacstd2*, and *Krt4,* adeno*–*L2) or basal (*Krt5* and *Krt14*, adeno*–*B) phenotypes. This conclusion was supported by bulk RNA–seq data from normal mouse L1, L2 and basal cells (Fig. 3A**, Fig. S7, refer to Methods**). While adenocarcinoma cells were readily annotated based on maximum likelihood (adeno*–*L1, *–*L2, and *–*B), these cells displayed significant basal and L2 phenotypic mixing (higher entropy) compared with wild–type cells (P<10^-16^, Fig. 3A, right bar plot). We further found that NEPC cells had the highest similarity to wild*–*type L2 cells among the wild*–*cell types (Fig. 3A**, refer to Methods**). In all, our scRNA-seq and spatial imaging data together implicate stem–like luminal (L2) cells and potentially basal–like state as the source of plasticity following deletion of *Pten/Rb1* or *Pten/Rb1/Trp53* that can give rise to at least 4 distinct lineages, namely NEPC, *Pou2f3–Gfp*, EMT–*Gfp*, and *Tff3*–*Gfp*. While we cannot exclude that these lineages arise from early activation and/or leakiness of *Pbsn*–*Cre*, notably with respect to EMT–*Gfp,* there is compelling evidence through lineage tracing using luminal *Nkx3.1–Cre* and *Krt8–Cre* mice, and our spatial IF showing PANCK and SYP staining that transformed luminal cells readily transdifferentiate into NEPC (*23*). Specifically, and consistent with our data, L2 (SCA–1+) cells have an increased propensity to transdifferentiate to neuroendocrine cells upon androgen ablation compared with L1 (SCA–1–)(*24*).

**Figure 3.**
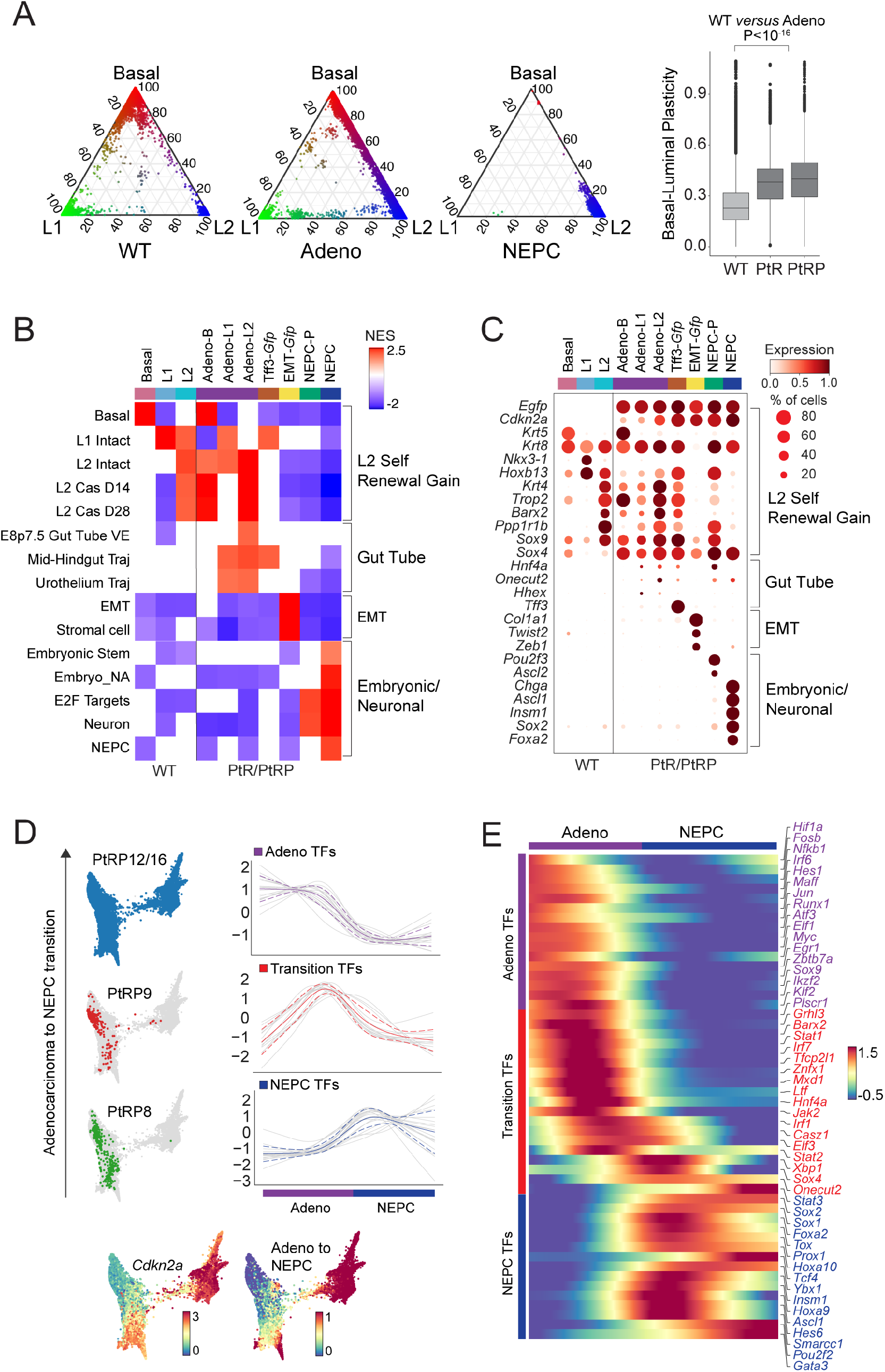
Stem– and Developmental–Like Programs Highest in L2–Like State with Mapping of Inferred Transition TFs. **(A)** Ternary plots of wild–type epithelial cells, adenocarcinoma or NEPC cells (left to right labelled ‘WT’, ‘Adeno’, and ‘NEPC’) based on three coordinates–basal (B), luminal 1 (L1), and luminal 2 (L2) Markov absorption probabilities. A preponderance of ‘Adeno’ cells (middle) along the B–L2 side of the triangle suggests an increased mixed phenotype compared with ‘WT’. Ternary plot of ‘NEPC’ cells (right) demonstrating skew to luminal 2 (L2) absorption probabilities. Box plot of basal–luminal plasticity *per* single cell, as measured by Shannon entropy of re–normalized Markov absorption probabilities for B, L1 and L2 phenotypes comparing WT *versus* mutant Adeno in PtR and PtR (Mann–Whitney U test comparing entropies of WT *versus* mutant Adeno in PtR/PtRP, **refer to Methods**). **(B)** Heatmap of significantly enriched gene sets *per* cell type (scale from -2 to +2.5 of GSEA normalized enrichment score, P<0.05, **refer to Methods**). **(C)** Dot plot of select differentially expressed genes (DEGs), stratified by cell type (mean normalized log_2_X+1 expression scaled from 0 to 1; size of dot represents percent of cells expressing the respective gene) grouped by L2, gut tube specification, EMT, and embryonic/neuronal genes. (**D)** FDL of mutant adenocarcinoma (restricted to PtR model, including adeno–B and adeno–L2) and NEPCs cells (total N=16,593 cells) colored by genotype and time (left). Shown below are FDLs colored by normalized log_2_X+1 expression of *Cdkn2a* or pseudotime scaled from 0 to 1 (bottom). Gene trends for TF DEGs across the adenocarcinoma to NEPC branch probability were determined as described in Palantir using a generalized additive model with cubic splines used as the smoothing function across 500 equally sized bins. Gene trends were grouped using Phenograph cluster (*k*=30). Solid and dashed lines represent the mean and standard deviation of the gene trends for each cluster (**Refer to Methods**). TFs groups include early adenocarcinoma (purple), transition (red), and NEPC (blue). (**E**) Heatmap of gene trends of select TF DEGs (Phenograph cluster versus rest, refer to **Methods: ‘Identifying DEGs’**) from each category are ordered by the adenocarcinoma to NEPC transition and colored by aforementioned groups (right; scale gene trends of imputed expression, -0.5 to 1.5).

Having documented a much greater than expected degree of lineage plasticity in human and mouse prostate cancers (i.e., beyond SYP+ NEPC alone), we studied transcriptional programs and transcription factors associated with these lineage transitions, starting with differentially expressed genes (DEG) and pathways using gene set enrichment analysis (GSEA) (**Tables S6, S7**). In addition to L2–like regenerative programs that we previously reported in the context of the normal castration response (*14*), lineage**–**specification programs of early gut development (*25, 26*) in *Gfp+* luminal and basal adenocarcinoma cells emerged (Fig. 3B **and** 3C). Among these programs, and specifically highest in adenocarcinoma luminal–like cells, we noted an upregulation of *Hnf4a* (focal positivity confirmed by IHC in **Fig. S6**), *Hhex* and *Onecut2,* which are known lineage**–**defining TFs in the developing foregut (*27, 28*). *Onecut2* is of particular interest due to its recently reported role in suppression of AR signaling in CRPC (*29*).

Given that the data suggest that reactivation of stem– and developmental–like programs in *Gfp*+ adenocarcinoma cells is the likely source of lineage plasticity, we looked more specifically at TFs that might be responsible, starting with a predefined set of known TFs (1637 genes, Transcription Factor Database, TFDB). Consistent with our FDLs (Fig. 2B), pseudotime analysis using Palantir (*30*) (**refer to Methods**) starting with an early adenocarcinoma state (cluster with low *Cdk2na* expression) revealed four terminal differentiated states: *Tff3*–*Gfp*, EMT–*Gfp*, *Pou2f3–Gfp* and NEPC (**Fig. S8**). To explore the set of perturbed TFs further and given its clinical relevance, we focused on the adenocarcinoma to NEPC transition, restricting to the PtRP model, and noted three major trends in TF expression, namely: (i) those active in adenocarcinoma alone, (ii) those expressed during the transition to NEPC and (iii) NEPC–specific regulators (Figs. 3D **and** 3E**, Fig. S8,** discussed later).

A closer look at the NEPC and non-neuroendocrine variant populations provided evidence for even more heterogeneity, consisting of three subsets defined by distinct DEG profiles designated as: (i) *Pou2f3–Gfp* (*Pou2f3, Dclk1*), (ii) NEPC–A (*Ascl1*), (iii) NEPC–Ar (**refer to Methods,** Fig. 4A**, Fig. S9**). Because the NEPC–Ar subset is largely based on scRNA-seq data from a single PtRP mouse, we confirmed the presence of NEPC–Ar populations by multiplex–IF in additional PtR and PtRP mice (**Fig. S9**). We further found a cluster of *Syp*+ cells that express both *Ar* and *Ascl1*. We asked if the diversity of NEPC lineages detected in the prostate GEMMs are also found in human CRPC (beyond that observed in our scRNA–seq cohort, Figs. 1 **and** 4A). To this end, we assessed expression of selected lineage markers across a cohort of tissue samples from a large rapid autopsy cohort (*31, 32*) containing tissue samples from multiple anatomically distinct metastatic sites *per* patient [16 patients, 139 tumors from distant metastases and the prostate gland; 2–21 metastatic sites per patient] (Fig. 4B, 4C, **Fig. S10, Table S8**). The analysis confirmed 2 of the 3 mouse NEPC subsets (NEPC–A and NEPC–Ar) in human CRPC. Human NEPC–A is notable for concordant INSM1 and DLL3 expression, consistent with the mouse profile, and reminiscent of ASCL1+ human SCLC. Human NEPC–AR samples were largely INSM1– and ASCL1–negative (Fig 4B). NEUROD1 and POU2F3 were absent in this subset at the TMA level.

**Figure 4.**
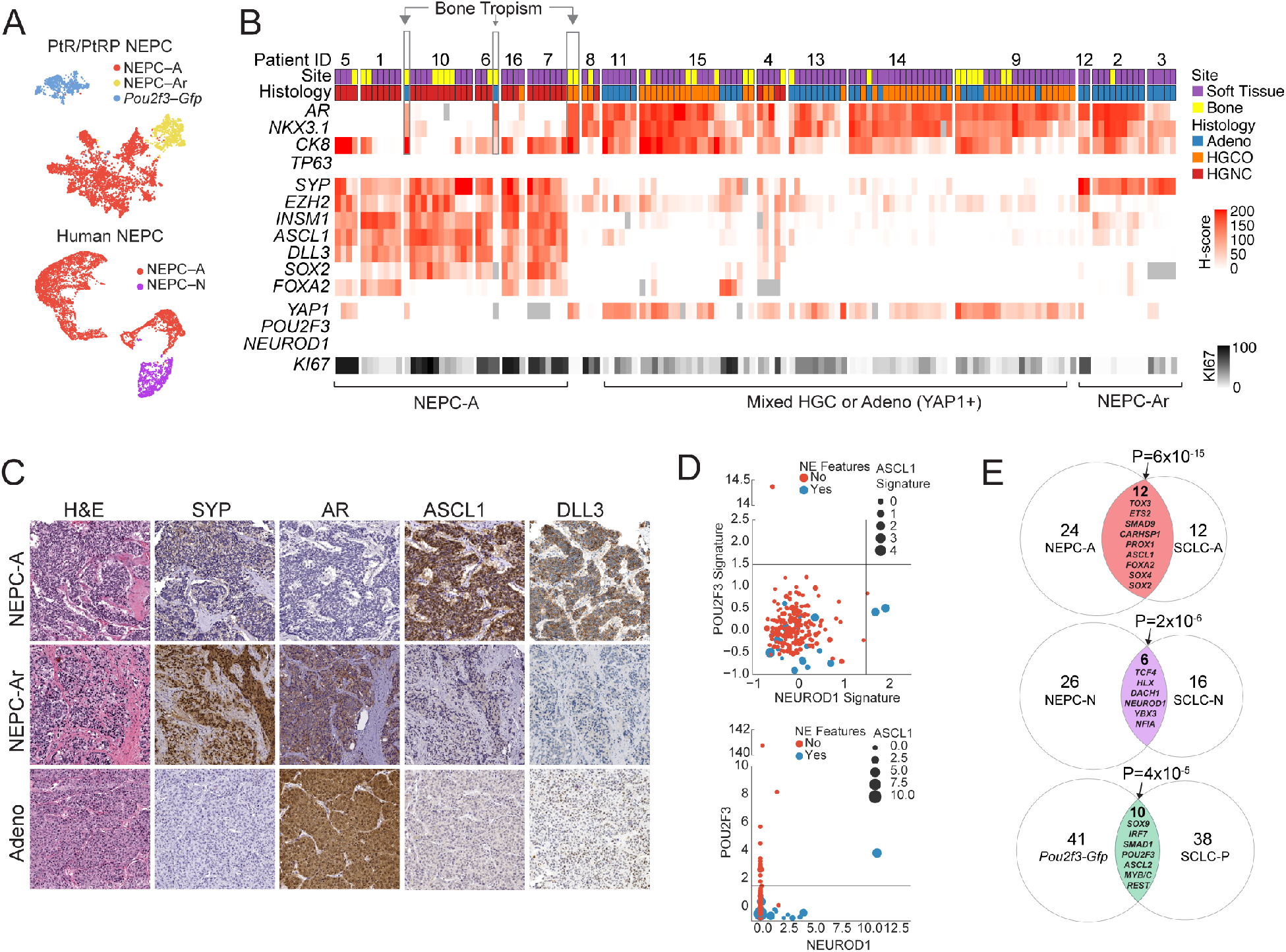
Conservation of Transcription Factors in NEPC Subtypes. **(A)** UMAP colored by NEPC subtypes based on *Ascl1* (A), *Ar* (Ar) and *Pou2f3* (P) in GEMMs (N=4,973 cells) (A) and *ASCL1* (A) and *NEUROD1* (N) in human samples (from HMP04, HMP16, and HMP17, N=9,572 cells). **(B)** Heatmap of human CRPC tissue microarray *H*-score (immunohistochemical score, scale 0 to 200, red gradient) and proliferative score (KI67, 0 to 100, black gradient) shown with corresponding patient ID, site (soft tissue, purple; bone, yellow), and histology (adenocarcinoma, blue; high-grade carcinoma, NOS, orange; high-grade neuroendocrine, red) confirming NEPC–A and NEPC–Ar subsets. Arrow denotes bone tropism of AR positivity in patients 6, 7 and 10. **(C)** Representative immunohistochemistry shown for NEPC–A, NEPC–Ar, and high-grade adenocarcinoma (additional examples and stain shown in **Fig. S10**). **(D)** Scatter plot shows *Z*–score of expression of genes in the SU2C database of *Pou2f3–Gfp* (y–axis) and NEPC-N (x–axis) signatures (top, **Refer to Methods**), and canonical transcriptional regulators *POU2F3* (y–axis) and *NEUROD1* (x–axis) (bottom). Red and blue dots denote non–NEPC or NEPC features as *per* SU2C database, respectively. Size of dot corresponds to the Z–score of NEPC–A signature (top) or ASCL1 expression (bottom). Dotted lines denote Z–score cutoff of 1.5. **(E)** Venn diagrams of differentially expressed TFs in corresponding *ASCL1*, *NEUROD1* and *POU2F3*–high subtypes in small cell lung cancer and neuroendocrine prostate cancer. The overlap between NEPC and SCLC subsets is assessed by Fisher’s exact test. Key overlapping TFs are annotated suggesting conserved transcriptional regulation.

Our TMA also revealed a subset of YAP1+ human high–grade carcinomas (HGC), while NEPC tumors largely showed absence or focal *YAP1* expression (*33*). Although the former finding may implicate altered Hippo pathway signaling in CRPC adenocarcinoma, important to note is that *YAP1* is also expressed in primary adenocarcinoma cells (*28*). Finally, when examining lineage marker expression across metastases, we found several examples of widespread SYP+ soft tissue metastases in patients whose bone lesions retained the expression of luminal epithelial markers (*AR*, *NKX3.1*, *CK8*), suggesting microenvironmental signals may influence lineage commitment (Fig. 4B, **Table S8,** patients 6,7, and 10, boxed in gray).

To determine whether there is a human counterpart to mouse *Pou2f3–Gfp*, we examined existing human CRPC (SU2C) and patient–derived xenograft datasets (PDXs, LuCAP) for expression of *POU2F3* and its signature (*12*) (**refer to Methods**). We also studied NEPC–N (*NEUROD1*) in both datasets; this subpopulation was found in patient HMP17 by scRNA–seq (Fig 1D), but not in mouse NEPC or by human TMA. Most NEPC samples in the SU2C dataset expressed *ASCL1,* as expected, but we did identify several *NEUROD1*–expressing samples (NEPC–N+) that also retained expression of ASCL1, in line with HMP17 and recent literature (*34*) (Z**–**score > 2, discussed above) (Fig. 4D). *Pou2f3–Gfp* was less convincing in that the extreme *POU2F3* outlier (by signature) is from a subcutaneous tumor, which may alternatively represent contamination from skin cells expressing high levels of POU2F3 (Fig. 4D). Similar to our GEMM data, analysis of the PDX dataset also revealed divergent NEPC–A and *Pou2f3–Gfp* phenotypes in two sublines derived from the same patient, LuCAP173. Notably, subline LuCAP173_2 (DNPC with rare foci of NEPC) showed high *POU2F3* expression, whereas subline LuCAP173.1 (fully NEPC) expressed low *POU2F3*, but high *INSM1, CHGA,* and *SYP* (**Fig. S11**). This is consistent with a recent cohort of human CRPC patients displaying some degree of POU2F3–positivity (*12*). While these findings are suggestive of the presence of these NEPC subsets, large scale tissue**–**based PDX and tumor IHC/IF are likely needed to delineate their prevalence. Lastly, we also found evidence of EMT–like programs in the SU2C dataset, although not exclusively linked to NEPC histology (**Fig. S11**). This observation was consistent with EMT programs being active in late stage CRPC, such as in our human single–cell analysis and enriched in HMP08 (DNPC, Fig. 1E). Furthermore, while *TFF3* was expressed in SU2C samples (highest *Z*–score of ∼8), the *Tff3–Gfp* signature for all samples fell below a *Z*–score of 1.5, perhaps because *TFF3* defines an early population not enriched in CRPC patients (as seen in the GEMM data, Fig 2D**, Fig. S11**).

Given the concordance of NEPC populations in the GEMM and human datasets (scRNA–seq, UW rapid autopsy, SU2C, and PDX), we asked whether the TFs that define these subsets are conserved across human SCLC subtypes. We found significant overlap of differentially expressed TFs in scRNA–seq data from human SCLC as well as in mouse and human NEPC subtypes (*35*) (Fig. 4E**, Fig. S7E**). Examples of shared NEPC and SCLC TFs include: *FOXA2, INSM1*, *SOX2*, *SOX4, TOX3* (ASCL1 subtype, P=6×10^-15^, Fisher’s exact test); *ASCL2*, *REST*, *IRF7, SOX9*, *MYC, MYB* (POU2F3 subtype, P=4×10^-5^, Fisher’s exact test) and *DACH1*, *NFIA*, *HLX*, *TCF4* (NEUROD1 subtype, P=3×10^-6^, Fisher’s exact test) **(Fig. S9**, additional overlap analyses). Collectively, these mouse/human and prostate/lung cancer pairs reveal a high level of conservation of the endoderm–derived lineage specification pathways, in essence, revealing how these can be hijacked to promote disease progression and drug resistance.

Because transition to NEPC portends a highly aggressive and lethal stage of prostate cancer, there is an acute need for novel therapies. Current drug development efforts are focused on actionable targets known to be expressed in NEPC, such as *EZH2*, *AURKA* and *DLL3*, with clinical trials currently underway. In light of the high degree of heterogeneity in NEPC seen through our single**–**cell analyses, we looked at expression of these and other common targets, such as *AR* and *PSMA* across our datasets. Not surprisingly, this analysis revealed a high degree of both intra–tumoral and inter–tumoral heterogeneity across both human and GEMM scRNA-seq (examples shown in Fig. 5A), raising obvious concerns about the clinical efficacy of these treatment strategies due to escape of target**–**negative tumor cells.

**Figure 5.**
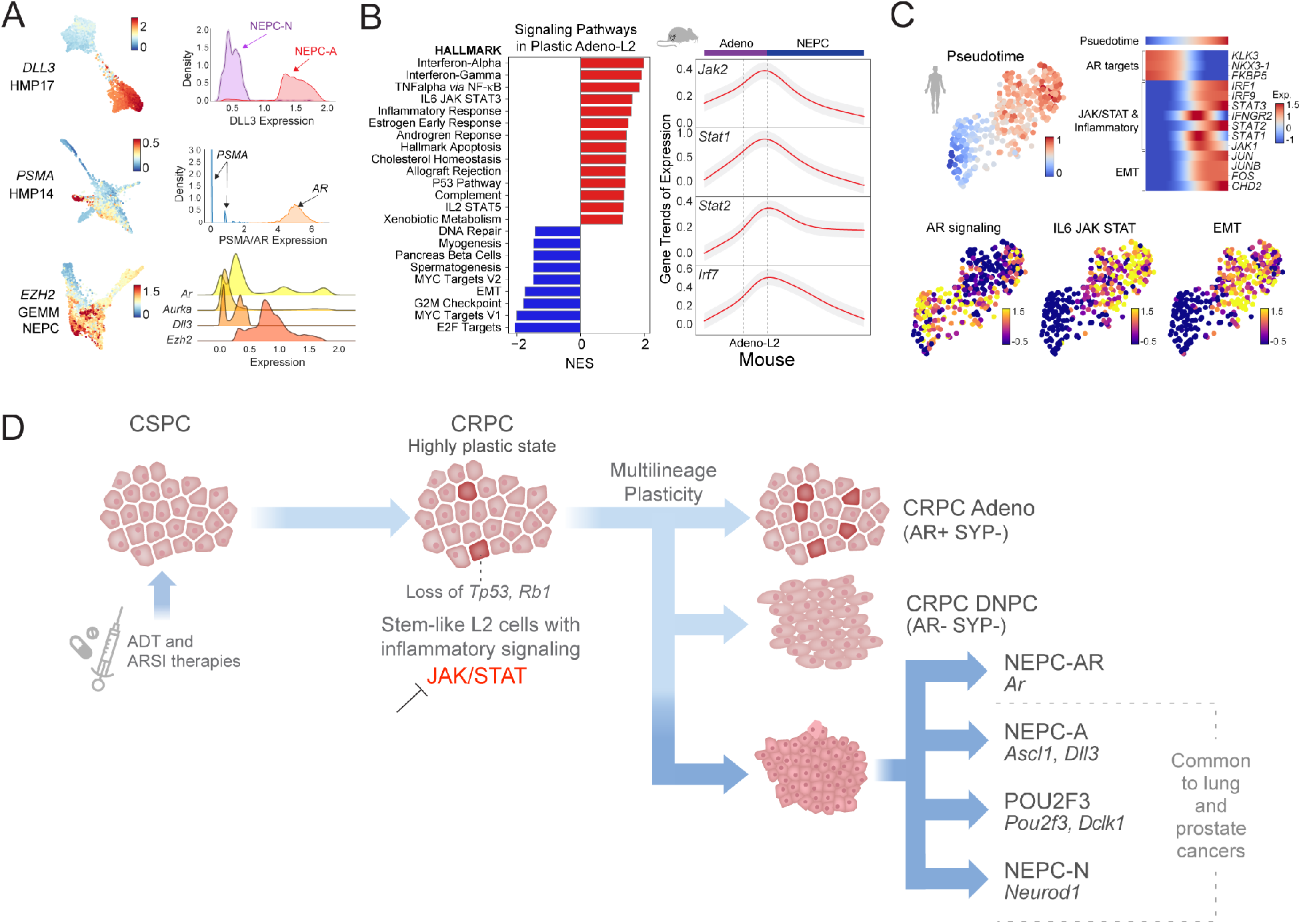
Identification of Targetable Transcriptional Mediators of Plasticity Across Human and GEMM Datasets. **(A)** FDL and kernel density estimate plots of imputed gene expression (MAGIC, *k*=30, t=3) are shown of current and select prostate cancer drug targets. FDL of *DLL3* expression is shown for HMP17 (NEPC, 2,845 cells, top) with density of expression stratified by NEPC–A (purple) and NEPC–N (red). FDL of *PSMA* expression is shown for HMP14 (CRPC adenocarcinoma, 3,351 cells, middle) highlighting both low *PSMA* (blue) and high *AR* (orange) expression. FDL of *Ezh2* expression is shown for GEMM NEPC tumors (N=4,973 cells, bottom) with ridgeline plots of imputed expression shown for *Ar, Aurka, Dll3, and Ezh2*. **(B)** Enriched signaling pathways using GSEA in adeno–L2 state highlighting inflammation and JAK/STAT pathways (scale -2.5 to 2, normalized enrichment score, P<0.05). Plotted gene trends of *Jak2*, *Stat1*, *Stat2, Irf7* were determined as described in Palantir across adenocarcinoma (purple bar) to NEPC (blue bar) using a generalized additive model with cubic splines used as the smoothing function across 500 divided equally sized bins. **(C)** UMAP of HMP13 (N=353 cells) colored by pseudotime from cells with high to low AR signaling (**refer to Methods**, top left). Corresponding robustness analysis of pseudotime detailed in **Fig. S11** (**refer to Methods**). Heatmaps showing gene trends for select DEGs and mouse transition TF markers ordered by the high to low AR signaling—early NEPC transition (right; scale -1 to 1.5, **refer to Fig. S11**). Gene trends were determined as described in Palantir and above in (B). AR signature score (**refer to Methods**) or Z–score of leading edge of enriched GSEA terms IL6–JAK–STAT or EMT are shown (scale, -0.5 to 1.5) (bottom). **(D)** Schematic of lineage plasticity progression and its mediators highlighting plastic L2–like cells enriched for *TP53, RB1, and/or PTEN* loss with emergence of developmental and stem–like program and upregulation of Jak/Stat and interferon signaling genes (ISGs). This cellular state is primed for multi–lineage plasticity with inhibition of Jak/Stat re–sensitizing to AR directed therapy (refer to accompanying manuscript). NEPC and neuroendocrine variant cells states, *ASCL1*, *NEUROD1*, and *POU2F3*, are common to both lung and prostate cancers with conservation of TFs. The identification of stem–like transformed L2 state provides a target for more focused therapeutic intervention.

Our prostate GEMM time course analysis implicated stem–like luminal (L2) adenocarcinoma cells as the source of plasticity for NEPC, raising the possibility that pharmacologic targeting of this population may be a strategy to prevent NEPC transition. We thus explored the temporal activation of pathways across the adenocarcinoma to NEPC transition in our GEMMs, with an eye toward druggability. In the adeno–L2, pathway analysis revealed the majority of significantly enriched gene sets related to activation of Jak/Stat, IFN and other inflammatory response pathways (Fig 5B). The set of genes (GSEA leading edge) driving this enrichment overlapped with transition regulators, namely IRF1/7, JAK2, STAT1/2, showing the highest expression in the adeno–L2 high plastic state, but with reduced expression in the NEPC populations (**Fig. S8**). To assess human relevance, we explored whether these transcription drivers were activated in tumor biopsies displaying evidence of plasticity from AR**–**high to AR**–**low adenocarcinoma states. Strikingly, we found a similar enrichment of inflammatory (and EMT) pathways in humans across high to low AR signaling as seen in HMP13 and highest in cells displaying an L2–like state (Fig 5C**, Fig. S11)**. Mapping of TFs across diffusion pseudotime revealed a similar activation of transcriptional regulators, namely JAK1, STAT2/3, IRF1, and IRF7 (Fig 5C**, Fig. S11**).

Together, our data reveal a greater than expected degree of lineage plasticity in prostate cancers that develop resistance to androgen deprivation therapy and, through side–by–side analysis of human and mouse datasets, establish a set of transcriptional programs that define and likely mediate plasticity states in prostate cancer. The delineation of such novel signatures should inform clinical prognosis and could enhance precision in treatment selection. Most importantly, we define a stem–like luminal (L2) population as the likely source of plasticity and implicate the inflammatory JAK/STAT pathway as a potential target for therapeutic intervention (Fig. 5D), which we address functionally in the accompanying manuscript.

## Acknowledgments

Authors are thankful to the members of the Pe’er and Sawyers laboratories for their extensive and productive critiques and discussions. We are incredibly grateful to the prostate cancer patients who participated in this research. We further appreciate the efforts of the Memorial Sloan Kettering Cancer Center (MSK) Genitourinary faculty for recruitment of prostate cancer tumor specimens. We thank the MSK core facilities for their invaluable help, namely the Molecular Cytology Core and Pathology Core for their help with confocal microscopy and IHC, and the Flow Cytometry Core for their help with FACS experiments. We are appreciative of the generous help of Sonja Nowotschin, PhD to curate a list of canonical gut tube specification transcription factors. We are similarly thankful to Matan Hofree, PhD for providing us with certain sets of developmental DEGs from published single cell data.

## Author contributions

S.Z., J.L.Z., J.M.C., D.P., and C.L.S. conceived the project. S.Z., J.L.Z., J.M.C., D.P., and C.L.S. wrote the manuscript. H.I.S, D.E.R, and M.J.M provided human tumor specimens. A.G., M.R. and M.F. interpreted histology and staining for prostate cancer patients. S.Z., J.L.Z., K.L., W.R.K and P.W. performed human tumor tissue dissociation. S.Z, J.L.Z., A.B. performed mouse staining and/or confocal microscopy. J.L.Z., K.W., K.L. performed all mouse work. M.P.R., N.R., I.L, J.S, A.J, and A.G. performed tissue microarray studies. P.S.N., C.M. and M.P.R. enrolled patients into rapid autopsy protocols, provided rapid autopsy biospecimens and RNA–seq data from PDX tumors. M.P.R. and M.C.H. interpreted and scored human tissue microarray. O.C., I.M., and T.X. performed single-cell sequencing. L.M., R.C. and D.P. oversaw the single-cell–sequencing experiments. S.Z. J.M.C., and J.C. performed computational analyses. D.P. and C.L.S. oversaw the project.

## Data and Material Availability

Human raw data for CSPC samples, as per *Karthaus et al*., are available at the Data Use and Oversight System controlled access repository https://duos.broadinstitute.org/ (accession no. DUOS-000115, samples: HP95NT/T, HP96NT/T, HP97NT/T, HP98NT, HP99NT/T, HP100NT/T, and HP101NT/T). Human raw data and 10X formatted files for CRPC samples are available at dbGAP (study submitted, accession no. in process). GEMM raw data and 10X formatted files for WT, PtR and PtRP are available at Gene Expression Omnibus respository (accession no. in process). Software and tools used for the enclosed data analysis will be provided open source at https://github.com/dpeerlab/.

## Supplementary Materials

Material and Methods

Figure S1. Clinical, molecular, and pathological features of castrate–sensitive (CSPC) and castrate–resistant (CRPC) prostate cancer biopsies.

Figure S2. Schematic, coarse cell typing, entropy and tumor annotation of human prostate cancer biopsies.

Figure S3. Intra–tumoral transcriptomic programs in select CSPC and CRPC adenocarcinoma biopsies.

Figure S4. Intra–tumoral transcriptomic programs in select DNPC and NEPC biopsies.

Figure S5. Schematic, coarse cell typing replicates, and entropy of genetically engineered mouse models (GEMMs).

Figure S6. Spatial immunofluorescence in GEMM time course and confirmation of *Tff3–, Vim–, Pou2f3–,* and *Hnf4A–*positive cells.

Figure S7. Phenotypic deconvolution of mutant cells in GEMMs.

Figure S8. Mapping transitions in adenocarcinoma to multi–lineage cell states in GEMMs.

Figure S9. Identification, characterization, and differential expression in NEPC subsets.

Figure S10. Extended select immunohistochemistry from human TMA data.

Figure S11. Mapping of human TFs from AR–high to AR–low–early–NEPC states and evaluation of GEMM cell types in PDX and SU2C datasets.

Table S1. Human and GEMM SEQC and scRNA–seq QC statistics.

Table S2. Clinical characteristics and molecular genetics of prostate cancer biopsies.

Table S3. Human and GEMM detailed antibody information.

Table S4. GSEA terms used in analysis (GMT format).

Table S5. DEGs by MAST of human prostate cancer subtypes.

Table S6. Enriched GSEA pathways in different GEMM cell states.

Table S7. DEGs by MAST in different GEMM WT, adenocarcinoma, and NEPC cell types.

Table S8. Complete TMA identifiers and H-scores per gene for TMA.

**References 36-57 only used in materials and methods section.

## Supplementary Materials

### Materials and Methods

#### Patient and GEMM cohorts

##### Human patient derived tissue

Tumor tissues were collected from patients undergoing a surgical resection or tissue biopsy for clinical care at Memorial Sloan Kettering Cancer Center (MSKCC). Informed consent was obtained for all patients and approved by MSKCC’s Institutional Review Board (IRB) #12–245 (NCT: 01775072), #06–107, and #12–001. Tumor biopsies included 6 primary castrate–sensitive prostate cancer (CSPC–HP) (*19*) and 10 metastatic castrate–resistant prostate cancer (CRPC–HMP). The 6 CSPC biopsies were previously analyzed and published by our group and are available at the Data Use and Oversight System controlled access repository: https://duos.broadinstitute.org/ (accession no. DUOS–000115, files annotated as HP95T to HP101T, all FASTQ files were re–processed *per* ‘**Data pre–processing’**) (*19*). All CRPC–HMP biopsies collected prospectively from 2019 to 2020 and were freshly derived from metastatic sites from patients with CRPC, who by definition, progressed after being treated with a next–generation anti–androgen therapy (ARSI) with or without a taxane. Serum PSA levels were reported from the date one week before or after of biopsy. Detailed clinical characteristics including AJCC staging, location, PSA, type and number of prior treatments, and genomic alterations are listed in **Table S2 and Fig. S1**. Routine clinical histology and immunohistochemistry were performed by a MSKCC pathologist. Based on this pathologic assessment, patients were classified into three clinically recognized phenotypes—namely CRPC adenocarcinoma (AR+, SYP–), double–negative prostate cancer (DNPC) (AR–, SYP–), or neuroendocrine with characteristic small cell features (AR–, SYP+) (*32*).

##### Targeted sequencing of human tumor tissue

For patients who consented to IRB 12–245 (NCT 01775072; MSK–IMPACT) and for clinical purposes, targeted sequencing was performed for a panel of actionable cancer genes for alterations (*36*) (**Table S2**). Genomic data was collected on retrospective review of the electronic medical record.

##### GEMM series

Mouse studies were carried out in compliance with Research Animal Resource Center guidelines (IACUC 06–07–012). The construction and genotyping of all GEMMs has been previously described (*9, 37, 38*). 7 PtR (7 mice, *PbCre*:*Rosa26*^mT/mG^*Pten*^fl/fl^*Rb1*^fl/fl^), and 13 PtRP (13 mice, *PbCre*: *Rosa26*^mT/mG^*Pten*^fl/fl^*Rb1*^fl/fl^*Tp53*^fl/fl^) mice analyzed were on a mixed C57BL/6:129Sv: FVB/NJ background. Timepoints for PtR included 24, 30, and 47 weeks, and for PtRP were 8, 9, 12, and 16 weeks relevant to the adenocarcinoma to neuroendocrine transition (*9, 37, 38*). Six PtRP mice were castrated at 8 weeks of age for either a total of 4 weeks (3 mice, ‘Cas’), or 2 weeks followed by 2 weeks of dihydrotestosterone (DHT, ‘Cas/Reg’) addback (3 mice). DHT pellet was implanted subcutaneously. Nine wildtype (WT) mice were on a matched or FVB background (Fig. 2A **and Fig. S5A**).

#### Sample Handling

##### Sample processing: human and GEMM tissue

Fresh benign and malignant tissue was mechanically cut using a scalpel into small pieces (∼1–5 mm^3^). The tissue was then processed and dissociated in 5–10 mg/ml collagenase type II (Gibco) solution in adDMEM/F12+/+/+ with 10 µM Y–27632 dihydrochloride for 30 minutes to 2 hours on a 37°C shaking platform. This was followed by a 1 minute 0.5 M EDTA wash at room temperature, and subsequent digestion with TrypLE (Gibco) with 10 µM Y–27632 dihydrochloride for 5–10 minutes at 37°C on a shaking platform until a single cell suspension was obtained (*39*). For human samples, if more than 10% of doublets were present (visual inspection) or there was evidence for <80% viability (hemocytometer, using 0.2% Trypan Blue), then cells were FACS–sorted for singlets and viability using a DAPI. For GEMM tissue, all cells were subsequently FACS–sorted for singlets and viability using DAPI.

##### Sample processing: 10X single cell RNA–sequencing

Dissociated cells were subjected to scRNA–seq using 10X genomics Chromium Single Cell 3’ Library and Gel bead Kit (v3 for human, except v2 for HP95T, and v2 for mouse) per manufacturer’s protocol. ∼3000 to 10,000 cells per sample was encapsulated and barcoded following the manual. Sample viability varied between 72 and 95% (0.2% Trypan Blue). The final sequencing libraries were double–size purified (0.6–0.8X) with SPRI beads and sequenced on Illumina Nova–Seq platform (R1–26 cycles, i7–8 cycles, R2–70 cycles or higher). For human samples, on average, 5861 cells per clinical biopsy were sequencing at a depth of ∼ 26,931 reads per cell (∼267 million reads per sample). The unique mapping was high, between 80.1–90.5%, with a median number of unique transcripts per cell being 4,973. For mouse samples, on average, 3,575 cells per mouse sample were sequenced at a depth of ∼ 48,685 reads per cell (∼213 million reads per sample). The unique mapping was high, between 73.4 – 80.4 %, with a median number of unique transcripts per cell being 4,403. Detailed QC statistics are listed in **Table S1**.

#### Immunohistochemistry, Immunofluorescence, and Tissue Microarray

##### Human immunohistochemistry of CRPC–HMP samples

For all CRPC–HMP biopsies AR or NKX3.1, SYP, and INSM1 stains were performed if not already completed for routine clinical work up **(Fig. S1)**. Exceptions included INSM1 stains for HMP16, HMP17 and SYP and INSM1 stains for HMP19 where additional tissue was not readily available. Briefly, and as part of routine clinical practice, patient biopsies were fixed in 10% neutral–buffered formalin, dehydrated with ethanol, and paraffin embedded per standard protocol. Furthermore, in select cases where ample formalin fixed tumor tissue (HMP04, HMP08, HMP11A/B, HMP13 and HMP14) was available, additional immunohistochemistry were performed by the MSK Clinical Pathology Core and include CK8, SYP, EZH2, POU2F3, ASCL1, INSM1, and YAP1. Immunohistochemistry of formalin–fixed tissue was performed using a Leica Bond RX and pretreated with epitope retrieval (ER1 or ER2) for 30–40 minutes. The respective primary antibodies were incubated for 30–60 minutes and followed by the Polymer Refine Detection Kit (Leica, DS9800) *per* the manufacturer protocol. Antibody clones, concentrations, antigen retrieval and platform used are listed in **Table S3**. Formalin–fixed stained tissue was scanned using a MIRAX scanner and processed using Case Viewer. All cases were reviewed by experienced genitourinary pathologists (A.G., M.R., M.C.H.) and immunoreactivities were scored in a blinded manner using a previously established H–score system, whereby the optical density level (“0” for no brown color, “1” for faint and fine brown chromogen deposition, and “2” for prominent chromogen deposition) was multiplied by the percentage of cells at each staining level, resulting in a total H–score (range 0–200) **(Fig. S1)** (*31, 32*).

##### Human tissue microarray

Tissue samples were procured from men who died of metastatic CRPC and who signed written informed consent to undergo a rapid autopsy as part of the Prostate Cancer Donor Program at the University of Washington as described previously (*31, 32*). The Institutional Review Boards of the University of Washington and of the Fred Hutchinson Cancer Research Center approved this project. In this study, a total of 139 anatomically distinct tumors including 47 adenocarcinomas, 54 high–grade carcinomas not otherwise specified (NOS), and 38 high–grade neuroendocrine carcinomas from 16 patients were analyzed. A TMA containing 2 separate cores for each metastatic site was constructed and adjacent sections were stained with antibodies specific to a total of 13 proteins including AR, NKX3.1, SYP, INSM1, ASCL1, SOX2, FOXA2, YAP1, CK8, DLL3, EZH2, TP63, NEUROD1, POU2F3, and KI67 (antibody specifications listed in **Table S3**). Tissue sections were counterstained with hematoxylin and slides were digitized on a Ventana DP 200 Slide Scanner (Roche). All cases were reviewed by experienced genitourinary pathologists (M.R., M.C.H.) and immunoreactivities were scored in a blinded manner using a previously established H–score system, whereby the optical density level (“0” for no brown color, “1” for faint and fine brown chromogen deposition, and “2” for prominent chromogen deposition) was multiplied by the percentage of cells at each staining level, resulting in a total H–score (range 0–200) (*31, 32*). The final H–score for each sample was the average of two replicate tissue cores. For KI67, any nuclear positivity was counted, and the percent of positive nuclei are shown (range 0–100). Included patient identifiers and H–scores for each protein are detailed in **Table S8**.

##### GEMM immunohistochemistry

Murine prostate tissue was fixed using 4% paraformaldehyde overnight, dehydrated with ethanol, and paraffin embedded per standard protocol. Immunohistochemistry (IHC) and immunofluorescence of FFPE tissue were performed using a Ventana Roche Benchmark Ultra (TFF3, rabbit polyclonal, Abcam, ab108599, 0.5 ug/mL) or Leica (HNF4A, rabbit monoclonal, Sigma Aldrich, ZRB1457, 0.06 ug/mL; and POU2F3, rabbit polyclonal, Sigma Aldrich, HPA–019652, 0.35 ug/mL) **(Fig. S6C)**. For TFF3 IHC, tissue sections were first heated for 32 minutes, followed by CC1 (Cell Conditioning 1, Ventana 950–500) retrieval and blocking using the Innovex (NB306) reagent for 30 minutes. The primary TFF3 antibody was incubated for 4 hours, followed by 60 minutes of incubation with biotinylated goat anti–rabbit IgG (Vector labs, PK6101) at 5.75 ug/mL. Blocker D, Streptavidin–HRP and DABMAP kit from Ventana Roche were used according to the manufacturer instructions. The slides were counterstained with hematoxylin and cover–slipped with Permount (Fischer Scientific). For HNF4A and POU2F3 IHC, tissue sections were first heated for 1 hour, loaded into the Leica Bond RX, and pretreated with an EDTA–based epitope retrieval ER2 solution (Leica, AR9640) for 20 minutes at 95°C. The respective primary antibodies (POU2F3 and HNF4A) were incubated for 60 minutes and processed with Polymer Refine Detection Kit (Leica, DS9800) *per* the manufacturer protocol. The slides were counterstained with hematoxylin and cover–slipped with Permount (Fischer Scientific). All FFPE stained tissue was scanned using a MIRAX scanner and processed using Case Viewer. All antibody concentration and antigen retrieval in **Table S3**.

##### Spatial immunofluorescence for GEMM

For quadruple spatial IF of GFP, VIM, E–CAD (Figs 2 **and S6**) and SYP, tissue sections were heated for 32 minutes, followed by CC1 (Cell Conditioning 1, Ventana 950–500) retrieval, and blocking using the Innovex (NB306) reagent for 30 minutes. First, a chicken polyclonal anti–GFP antibody (Abcam, ab13970) was used at 2 ug/ml. The incubation with the primary antibody was performed for 5 hours, followed by a 60–minute incubation step with biotinylated goat anti–chicken IgG (Vector labs, T1008) at 7.5 ug/mL. Blocker D, Streptavidin–HRP and TSA Alexa 488 (Life Tech, B40932) were then used for 16 minutes. Second, a mouse monoclonal anti–Vimentin antibody (Vector Lab, VP–V684) was used at 0.1 ug/mL. The incubation with the primary antibody was performed for 5 hours followed by biotinylated anti–mouse secondary (Vector Labs, MOM Kit BMK–2202) at 5.75ug/mL. Blocker D, Streptavidin–HRP and Tyramide–CF594 (Biotium, 92174) were used for 16 minutes. Third, a mouse monoclonal anti–E–cadherin antibody (BD Bioscience, 610181) was used at 2.5 ug/ml. The incubation with the primary antibody was performed for 5 hours followed by biotinylated goat anti–mouse secondary (Vector Labs, MOM Kit BMK–2202) at 5.75ug/mL. Blocker D, Streptavidin–HRP and TSA Alexa 647 (Life Tech, B40958) was used for 16 minutes. Fourth, a rabbit monoclonal anti–Synaptophysin antibody (clone YE269) (Epitomics, 1485–1) was used at 0.05 ug/ml. The incubation with the primary antibody was performed for 5 hours, followed by 60 minutes incubation with biotinylated goat anti–rabbit IgG (Vector labs, PK6101) at 5.75ug/ml. Blocker D, Streptavidin–HRP and CF 543 (Biotium, 92172) were used for 16 minutes. All slides were counterstained in 5ug/mL DAPI [dihydrochloride (2–(4–Amidino Phenyl)–6–indolecarbamidine dihydrochloride] Sigma D9542 for 5 minutes at room temperature, mounted with anti–fade mounting medium Mowiol [Mowiol 4–88] and cover–slipped. For quadruple spatial IF of GFP, SOX2, AR and SYP, (Figs. 2E **and S6**) the procedure was the same as above, but rabbit polyclonal anti–Sox2 antibody, (Chemicon–Millipore, AB5603) was used at 2 ug/mL in place of VIM (Step 2), and rabbit monoclonal anti–androgen receptor (AR) antibody (Epitomics, 3184–1) was used at 0.66 ug/mL in place on ECAD (Step 3). All FFPE stained tissue was scanned using a MIRAX scanner and processed using Case Viewer. All antibody concentration and antigen retrieval in **Table S3**.

#### Pre–processing of scRNA–seq data

##### Data pre–processing

For each sample, fastq files were processed individually using the SEQC pipeline with default parameters used for the 10X single–cell 3’ library. For human samples, 6 primary CSPC–HP and 10 metastatic CRPC–HMP were mapped to the hg38 human genome reference. For GEMM samples, 9 WT, 7 PtR and 13 PtRP were mapped to the mm38 mouse genome reference with a custom *Gfp* mapping. The SEQC (*40*) pipeline performed read alignment, multi–mapping read resolution with cell barcode and UMI correction to generate a count matrix [cells x genes]. The pipeline further performed the following initial cell filtering steps: true cells were distinguished from empty droplets based on the cumulative distribution of total molecule counts; cells with a high fraction of mitochondrial molecules were filtered >20%; and cells with low library complexity expressing very few unique genes were filtered (3 standard deviations (SDs) below the mean). In addition, we used the EmptyDrops (*41*) package and performed additional filtering of empty droplets by retaining cells with FDR <0.01. Putative doublets were removed using the DoubletDetection package (*42*) . Genes that were expressed in > 10 cells were retained for further analysis.

##### Combined matrix and normalization

For human samples, the entire cohort of CSPC–HP and CRPC–HMP samples yielded a filtered count matrix of 76,929 cells x 22,672 genes, with a median of 8,866 molecules per cell and a median of 3,901 cells per sample (**Table S1**). For GEMM samples, the entire cohort of WT, PtR, and PTRP tissues yielded a filtered count matrix of 67,622 cells x 19,591 genes, with a median of 8,626 molecules per cell and a median of 2,328 cells per sample (**Table S1**). For both human and GEMM datasets, the count matrix was normalized using sctransform (*43*) to obtain Pearson residuals per gene that adjust for technical artifacts in normalization. To visualize gene expression, non–zero UMI counts were corrected using sctransform (*43*), normalized by library size, scaled by median library size, and log2–transformed with a pseudo–count of 1.

#### Batch Correction

##### Summary of batch correction

For the human dataset, we used batch correction at the global level, propagating batch–corrected counts for downstream subsetting. For the GEMM datasets, batch correction was not required. Both batch–corrected human and non–batch–corrected GEMM datasets resulted in high coherence and mixing among immune populations, while maintaining distinct tumor phenotypes. For the GEMM dataset, replicates showed a high degree of mixing. Our methods, rationale, and evidence for batch correction for each dataset are noted below.

##### Batch correction of the human dataset

We performed batch correction of the entire human dataset including primary prostate samples (CSPC: HP tumors) and metastatic CRPC samples (HMP tumors) using fastMNN (*44*) with cosine distance applied to the residuals of 5,000 highly variable genes (minus cell cycle [regev_lab_cell_cycle_genes.txt] dataset, ribosomal and mitochondrial genes), reduced to the top 50 PCs with 40% of variance explained. We opted for fastMNN due to the ability to perform hierarchical merging among samples from the same patient (HMP11A and HMP11B).

##### Entropy–based measure of batch correction

To evaluate the effects of batch correction, we used an entropy–based measure that quantifies the degree of mixing of normalized expression across patients or GEMM samples (*40, 45*). We constructed a k–nearest neighbors (*knn*) graph (*k*=30) from the normalized matrix using Euclidean distance and computed the fraction of cells *q_T_* derived from each tumor sample *T* in the neighborhood of each cell *j.* We calculated the Shannon entropy *H_j_* of sample frequencies within each cell’s neighborhood as:

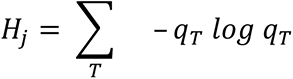

High entropy indicates that most similar cells come from a well–mixed set of tumors. For human datasets, and with batch correction, while many tumor cells remained distinct and patient–specific, there was high coherence and mixing among immune populations indicating effective batch correction (**Figs. S2B and S2C**). For GEMM datasets, there was high coherence of mixing the immune populations without batch correction (**Figs. S5C, S5D, and S5E**).

#### Measuring Inter–Patient Heterogeneity Per Cell Type

We used an entropy–based measure of inter–patient diversity for each tumor type, including benign prostate, CSPC, CRPC adenocarcinoma, DNPC and NEPC. We used the Phenograph clusters (*k*=30) within subsetted tumor cells (N=29,373 cells), myeloid, lymphoid, and mesenchymal populations without batch correction. Each cluster *C* represents a discrete phenotype of a given cell. To account for any differences in the number of cells per cluster and cell type, we subsampled 100 cells from each cluster 100 times with replacement and calculated the Shannon entropy of tumor biopsy frequencies *P* in each subsample *H_c_* as:

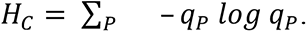

We then compared the distribution of Shannon entropies bootstrapped from clusters using a Bonferroni–adjusted two–sample t–test (Fig. 1C **and Fig. S2C)**.

#### Visualization of ScRNA–Seq Data

##### Visualization of different cell type compartments

To visualize all cells, or subsets of cell types (epithelial, mesenchymal, lymphoid and/or myeloid, individual HMP samples, or *Gfp*–positive cells), we used either uniform manifold approximation and projection (UMAP) (*46*) or force–directed layout (FDL) constructions (*47*). UMAPs were generated by creating a partition–based graph abstraction (PAGA) based on Phenograph (*21*) clusters that served to initialize the low dimensional embedding. The effective minimum distance between embedded points was set to 0.2 to 0.3 (*knn*=30, min_dist=0.2–0.3, and init_pos=‘paga’). To visualize cell state transitions and local relationships, FDLs were constructed. First, a *knn* graph was created based on PCs. The corresponding adjacency matrix was converted to an affinity matrix using an anisotropic Gaussian Kernel with *k* ranging from 20 to 40 neighbors as the scaling factor. The affinity matrix was then used as input for the draw_graph scanpy function––an implementation of the ForceAtlas2 (*47*) python module––to compute the FDL. Given the sparse nature of single–cell sequencing that arises from gene dropout, for certain analyses which we have explicitly commented on in the Methods, we imputed gene expression using MAGIC (k= 30, t=3) (*48*).

#### Phenograph Clustering

##### Parameter selection and robustness analysis

We performed unsupervised clustering of single cells using Phenograph (*21*). To calculate robust clusters, we performed Phenograph clustering over a range of values for the parameter *k* (numbers of neighbors in the *knn*–graph) to ensure that cell typing, and cluster labelling was consistent. We calculated the adjusted Rand index to determine the consistency of clustering across different *k* (10 to 80). The optimal *k* = 30 for all cell compartments was chosen from the window where the adjusted Rand index was consistently highest.

#### Human Cell Type Annotation

A schematic of human cell typing and subsetting is shown in **Fig. S2A.**

##### Human coarse cell type identification and subsetting

We used a hierarchical strategy to identify cell types using expression of canonical marker genes in unsupervised clusters. We first identified coarse cell types by performing Phenograph clustering (*21*) (*k*=30) on the batch–corrected count matrix (refer to **“Batch correction of human dataset”**). This step yielded 43 clusters spanning 76,929 cells. Clusters were annotated by expression of canonical markers, including *EPCAM* or *CHGA/CHGB* (epithelial, 44,448 cells), *PTPRC* and either *CD2/CD3E*, *CD79A*, or *TPSB2* (lymphoid; 11,152 cells), *PTPRC* and *CD14* and/or *LYZ* (myeloid; 6,121 cells), *COL1A1/VIM* and/or *CLDN5* (mesenchymal and endothelial; 7,922 and 5,484 cells, respectively) (**Fig. S2B**). We also used a more comprehensive gene list to confirm our coarse cell types referencing PanglaoDB, https://panglaodb.se/). We subsetted each coarse cell type for downstream analyses *with* propagation of batch correction, as described in the following sections.

##### Human designation of epithelial and tumor cells

*EPCAM*–positive epithelial clusters (N=44,448 cells, which includes 19,250 primary cells from CSPC–HP series and 25,198 metastatic cells from CRPC–HMP series). We projected sctransform residuals (batch–corrected in the combined dataset) onto the top 29 principal components (PCs) detected by knee–point with 98.5% variance explained. We executed Phenograph clustering (*k*=30), yielding 45 clusters.

We obtained cells from primary CSPC–HP tumors that were previously annotated as benign vs malignant in *Karthaus* et al. (refer to Karthaus *et al.* Methods section **‘Identification of potentially malignant cells by inferred copy number alterations’**, provided kindly by Matan Hofree, PhD) (*2*). We further considered any basal cells detected in CSPC–HP tumors to be benign, as a hallmark of primary adenocarcinoma is the loss of basal (*TP63*–positive) cells. These steps left a total of 4,333 primary CSPC adenocarcinoma cells (classified as malignant and comprising 22% of total epithelial cells). To support our cancer cell identification, we assessed the expression of genes known to be upregulated in malignant prostate cancer, including *KLK3/PSA, PSMA/FOLH1, ETV1* (up in HP97)*, SOX9,* and *CDKN2A*. We noted that in contrast to benign basal and luminal cells, malignant CSPC–HP cells showed higher expression of these genes compared with benign cells (**Figs. S2D, S2E**).

To identify malignant cells in CRPC–HMP metastatic biopsies, we followed a similar marker–based approach and considered any cluster with expression of known oncogenic or prostate–specific markers (*AR* or *KLK3/PSA, CDKN2A, CHGA* or *CHGB*) in a metastatic site to be malignant (**Figs. S2D, S2E)**. We however did note that cluster 41 (from HMP05) had high expression of *ALB, APOA1, CRP* (hepatocyte markers) (refer to PanglaoDB, https://panglaodb.se/) without *CDKN2A* expression (upregulated with *TP53/RB1* loss as in HMP05), compared to cluster 22 from the same patient (HMP05) that had low expression of *ALB, APOA1, CRP,* but high expression of *CDKN2A.* We therefore concluded that cluster 41 is likely a benign hepatocyte cluster given that this biopsy was derived from a liver metastasis. Furthermore, we ensured that putative cancer subpopulations clustered separately from cells derived from the normal prostate.

As a parallel approach to confirm our identification of malignant cells in both CSPC and CRPC, we assessed single–cell CNV profiles, based on the reasoning that any cells with a high degree of CNVs are likely malignant. To identify CNV profiles, we used a sliding window approach executed in the infercnvpy package (*49*), using a sliding window of 100 genes, with diploid mean and standard deviation (SDs) defined by benign luminal cells annotated in *Karthaus et al* (*2*). We calculate a summary score of CNV burden per cluster based on the mean of the absolute value of CNV. As expected, we find that malignant cells show significantly higher CNV burden compared to primary benign cells (**Figs. S2F, S2G**, P<6×10^−78^, Student’s t–test). We further noted that the CNV score was highest in metastatic NEPC/DNPC samples, followed by CRPC adenocarcinoma, and then primary adenocarcinoma (CSPC) (**Figs. S2F, S2G**). Overall, our approach yielded a total of 29,373 malignant cells (composed of 25,040 CRPC–HMP cells and 4,333 CSPC–HP cells).

##### Human designation of adenocarcinoma and NEPC cells

Tumor designated cells as above (N=29,373 cells) were subsetted (**Fig. S2A)**. We projected sctransform residuals (batch–corrected in the combined dataset) onto the top 24 principal components (PCs) detected by knee–point with 98.5% variance explained. We identified neuroendocrine and non–neuroendocrine cell types by first implementing Phenograph clustering (*k*=30) yielding 34 clusters which we labelled to be neuroendocrine based on expression of canonical markers (*CHGA, CHGB, SYP, ASCL1, INSM1).* We thereafter separated neuroendocrine (14 clusters, 13,696 cells) and non–neuroendocrine (20 clusters, 15,677 cells) clusters. For neuroendocrine cells, we projected the normalized counts on the top 9 PCs detected by knee–point with 98.7% variance explained Phenograph clustering (k=30) yielding 23 clusters. We used well published markers for different small cell lung cancer subtypes (*45*), including *ASCL1, NEUROD1, YAP1,* and *POU2F3* to annotate clusters. We noted three major groups with ASCL1, NEUROD1, and CHGB–alone (the latter with no other canonical NE markers such as *SYP, CHGA, ASCL1, NEUROD1*) expression. We subsequently labelled these as NEPC–A (13 clusters, 7,368 cells), NEPC–N (4 clusters, 1,574 cells), NEPC–CHGB only (7 clusters, 4,754 cells). NEPC–CHGB cells were largely derived from HMP08, a sample histologically classified as a DNPC with predominant basaloid morphology, and which on our single cell analysis displayed both *CHGB*+ and *CHGB*–tumor cells **(Fig. S4A)**. For adenocarcinoma cells, we projected sctranform residuals (batch–corrected in the combined dataset) on the top 12 PCs detected by knee–point with 98.5% variance explained Phenograph clustering (k=30) yielding 27 clusters. All clusters showed expression of either *AR, KLK3/PSA* or *CDKN2A* suggestive of transformed metastatic prostate cancer.

##### Human characterization of CRPC samples

For each CSPC–HP and CRPC–HMP sample, we subsetted tumor cells, re–normalized counts, and projected the top PCs as selected by knee–point. We executed Phenograph clustering (*k*=30). These steps yielded the following for CSPC–HP samples: HP96 (355 cells, 86 PCs, 55% variance explained), HP97 (1726 cells, 116 PCs, 44% variance explained, **Fig. S3A**), HP99 (1296 cells, 107 PCs, 57% variance explained, **Fig. S3B**), HP100 (406 cells, 50 PCs, 49% variance explained), HP101 (550 cells, 138 PCs, 57% variance explained). These steps yielded the following for CRPC–HMP sample: HMP04 (1791 cells, 94 PCs, 40% variance explained, **Fig. S4B**), HMP05 (501 cells, 147 PCs, 65% variance explained), HMP08 (4279 cells, 88 PCs, 32% variance explained, **Fig. S4A**), HMP11A/B (5718 cells, PCs, variance explained), HMP13 (354 cells, 102 PCs, 52% variance explained, **Fig. S3C**), HMP14 (3351 cells, 88 PCs, 28% variance explained, **Fig. S3D**), HMP16 (4936 cells, 75 PCs, 28% variance explained), HMP17 (2845 cells, 123 PCs, 35% variance explained, **Fig. S4C**), HMP19 (1265 cells, 94 PCs, 53% variance explained). Differential expression and pathway analysis was performed for each sample (refer to **“Identifying DEGs and Enriched Pathways**”). Specific DEGs were plotted by UMAP for select biopsies to highlight intra–tumoral heterogeneity (**Fig. S3–S4, and** Fig. 2D). Patient level clusters largely recapitulated clusters identified at the cohort level of analysis. However, it is interesting to note that by combining patients, increasing the number of cells, and incorporating trends that were shared across patients we got clearer subtypes that explained a far larger percentage of the variance.

##### Human mesenchymal annotation

*COL1A1/COLA12* and *CLDN5*–positive mesenchymal clusters (N=14,787). We projected sctransform residuals (batch–corrected in the combined dataset) onto the top 19 principal components (PCs) detected by knee–point with 98.6% variance explained. We executed Phenograph clustering (*k*=30), yielding 25 clusters. Cells were annotated as fibroblasts (16 clusters, 9,660 cells, *COL1A1*, *COL1A2*, *COL6A2*, *MFAP4, PDGFRB*) or endothelial (10 clusters, 5,127 cells, *CLND5, PECAM1, FLT1, FLT4, VWF*).

##### Human lymphoid and myeloid annotation

We subsetted CD45+ myeloid (monocyte, macrophages, PMNs, and DCs, 6,121 cells, *CD14, LYZ, CD14, TPSAB1/TPSB2*). We projected sctransform residuals (batch–corrected in the combined dataset) onto the top 13 principal components (PCs) detected by knee–point with 98% variance explained. We executed Phenograph clustering (*k*=30) yielding 17 clusters. We identified monocyte/macrophages (4,332 cells, 12 cluster, *HLA–DRA, HLA–DRB1, LYZ*), polymorphonuclear neutrophils (525 cells, 1 cluster, *ITGAX, CSF3R*), dendritic (550 cells, 1 cluster, *CLEC10A*), and mast (714 cells, 4 cluster, *TPSAB1/TPSB2*) cells. We similarly subsetted CD45+ lymphoid (B cells, T cell, NK cells, 11,152 cells). We projected sctransform residuals (batch–corrected in the combined dataset) onto the top 13 principal components (PCs) detected by knee–point with 98.5% variance explained. We executed Phenograph clustering (*k*=30), yielding 19 clusters. We identified NK/T cells (*CD3E, CD8, CD4, NKG7, KLRD1, GZMB,* 16 clusters, 10,097 cells) and B cells (*CD79A, MS4A1,* 4 clusters, 1,055 cells).

#### GEMM Cell Type Annotation

A schematic of GEMM cell typing and subsetting is shown in **Fig. S5B.**

##### GEMM coarse cell type identification and subsetting

We used a hierarchical strategy to identify cell types (**Fig. S5B**). We identified coarse cell types by first implementing Phenograph clustering (*21*) (*k*=30) on the normalized count matrix (refer to **“Data Pre–Processing”**). This yielded 44 clusters spanning 67,622 cells **(Fig. S5C)**. Clusters were annotated by expression of canonical markers, which included *Epcam* or *Chga/Chgb* (epithelial, 29,380 cells), *Ptprc* and *Cd2/Cd3e* or *Cd79a* (lymphoid, 6,441 cells), *Ptprc* and *Cd14* and/or *Csfr1 or Csfr3* (myeloid, 13,491 cells), *Col1a1* and/or *Cldn5* (mesenchymal and endothelial, 6441 cells). We also used a more comprehensive gene list to confirm our coarse cell types referencing PanglaoDB, https://panglaodb.se/). We subsetted each respective cluster for downstream analysis *without* batch correction **(Fig. S5B)** – this was possible because batch effects were not observed (entropy panel of **Figs. S5C, and S5D/E**).

##### GEMM epithelial compartment

*Epcam*–positive epithelial clusters (N=29,380 cells) were subsetted. We projected sctranform residuals without batch correction onto the top 80 principal components (PCs) detected by knee–point with 63% variance explained. We executed Phenograph clustering (*k=30)*, yielding 39 clusters. Clusters expressing high levels of *PATE4* (seminal vesicle marker), *CALML3* (basal seminal vesicle marker), or *FOXI1* (L3 marker) (*2*) were removed (2,839 SV cells and 817 L3 cell removed) as this latter subset is highly represented in the epididymis (*2, 15*). Furthermore, L3 cells, even after removal of its epididymal subpopulation, have been shown by *Karthaus et al.* to have no significant transcriptomic changes during a cycle of castration and generation (*2*). After removal of these cells, a total of 25,724 epithelial cells remained. All mutant PtR and PtRP epithelial cells fell into *Gfp*–positive clusters, consistent with *a priori* expectation that Cre recombination would be highly efficient (*9*).

##### GEMM mesenchymal compartments

*Col1a1* or *Vim*–positive mesenchymal clusters (N=18,310 cells) were subsetted (**Fig. S5B)**. We projected sctranform residuals without batch correction onto the top 89 principal components (PCs) detected by knee–point with 44% variance explained. These steps yielded 39 clusters. As there were both *Gfp* positive and *Gfp* negative clusters, we used the following strategy to identify transformed *Gfp–*positive mesenchymal/EMT–like cells. We plotted the median imputed expression of *Gfp* per cluster, which showed a bimodal distribution. A mixture model of two Gaussians showed that a threshold of 1.6 separated the bimodal distribution of *Gfp* expression well. We therefore labelled clusters with a median imputed *Gfp* > 1.6 as EMT–*Gfp* (clusters 1, 12, 13, 14, 24, and 27). We further excluded 216 cells from wild–type (WT) mice that also were captured in these clusters. These steps yielded a total of 3,210 EMT–*Gfp* cells. Given the lack of clear separation between mesenchymal cells from WT mice and *Gfp* positive mesenchymal cells, we cannot exclude the possibility of recombination of PtR or PtRP in wild–type mesenchymal cells due to the *Pbsn* promoter.

##### GEMM wild–type epithelial annotation

To define the wild–type epithelial cellular architecture in from our 9 wild–type (WT) mice, we subsetted WT epithelial populations (from ‘**GEMM Epithelial compartment**’, 7,435 WT cells from total 29,380 epithelial cells), again excluding seminal vesicle (SV) and luminal 3 (L3) cells. We first projected sctranform residuals onto the first 70 PCs selected by knee–point, explaining 50% variance. We then executed Phenograph clustering (*k*=30) yielding 23 clusters. Based on previously published canonical markers (*2, 15, 50*), we labelled wild–type epithelial cells as basal (*Krt5*, *Krt14*; 5808 cells, 14 clusters), L1A (luminal 1 anterior lobe, *Nkx3.1*^high^, *Hoxb13*^low^; 523 cells, 3 clusters), L1D (luminal 1 dorsal lobe, *Nkx3.1*^high^, *Hoxb13*^high^; 345 cells, 3 clusters), L1L (luminal 1 lateral lobe, *Sbp*; 254 cells, 1 cluster), L1V (luminal 1 ventral lobe, *Msmb*^high^; 322 cells, 2 clusters), and L2 (luminal 2, *Krt4*, *Tasctd2, Sca1*; 183 cells, 1 cluster) (**Fig. S7B**).

##### GEMM mutant*–gfp* cells PtR and PtRP annotation

To understand the different cell types in the mutant *Gfp*–positive populations, we subsetted all mutant epithelial and *Gfp–*positive mesenchymal cells (‘**GEMM Epithelial compartment**’ without SV and L3 cells and restricting to PtR and PtRP genotypes, and **‘GEMM Mesenchymal compartment’** with *Gfp*–positive *cells*). These steps yielded 21,499 cells. We first projected sctranform residuals onto the first 89 PCs selected by knee–point, explaining 64% variance. We then executed Phenograph clustering (*k*=30) yielding 31 clusters, and labelled clusters based on expression. We noted four groups of cells: (1) a large *Epcam*– and *Gfp*–positive population (18 clusters), (2 and 3) two additional smaller *Epcam*– and *Gfp*–positive population (*TFF3,* 1 cluster; and *POU2F3* and *ASCL2,* 3 clusters), and (4) a *CHGA–*positive neuroendocrine population (*CHGA, CHGB, SYP,* 6 clusters), and (5) a EMT–like population (*VIM, COL1A1, NCAM1,* 3 clusters). We labelled these as (1) Adeno (adeno–*Gfp*), (2) *Pou2f3–Gfp* (*POU2F3, ASCL2),* (3) *Tff3*–*Gfp,* (4) EMT–Gfp, and (5) NEPC, respectively (Fig. 2C **and** 2D**, Fig. S5C and S7C)**. In order to further delineate sub adenocarcinoma–*Gfp* populations, we trained a classifier based on wild–type cells as a training set (refer to **‘Phenotype deconvolution of GEMM *Gfp*–positive mutant cells’**).

##### Pooled dataset of GEMM WT epithelial and GFP+ populations

To understand the changes and potential trajectories from wild–type epithelial cells to *Gfp–*positive mutant cells, we subsetted both WT epithelial (**“GEMM WT epithelial annotation”** 7,435 cells) and mutant *Gfp–*cells (“**GEMM Mutant*–Gfp* Cells PtR and PtRP mice**” 21,499), yielding a total of 28,934 cells. PCA was performed with the top 76 principal components (PCs) retained with 62% variance explained. FDL are shown in Figs. 2C, 2D**, and S7C** (refer to “**Visualization of scRNA–seq data”**).

##### Phenotype deconvolution of GEMM *Gfp*–positive mutant cells

To deconvolute the mutant *Gfp+* cell types and assess the degree of cell type mixing in adeno–*Gfp,* we performed cell type classification of the mutant *Gfp*–positive populations by calculating Markov absorption, as executed in the Phenograph package (*21*), which can model mixed cancer phenotypes (*51*) (**Fig. S7A)**. For this approach, we combined mutant *GFP*–positive cells with WT epithelial cells (as described in “**Pooled dataset of GEMM WT epithelial and GFP+ populations”**, N=28,934 cells) and used the wildtype prostate epithelial cells (refer to **“GEMM WT epithelial annotation”**) as training data (L1A, L1D, L1V, L1L, L2, and B1) with annotated cell types (**Fig. S7B)**. Markov absorption models the probability that each unlabeled mutant cell represents a given cell type based on random walks on a graph. Each mutant cell can therefore be described by a simplex of cell–type probabilities summing to 1. For each mutant cell, we can then assign a hard classification of cell type by maximum likelihood or model the mutant cell as a mixed phenotype (*45*).

To ensure that wild–type cells were equally represented as the training data, we down–sampled to 183 WT cells per category, which included L1A, L1D, L1V, L1L, L2, and B1 (**Fig. S7B,** lowest number of cells is L2 is 183 cells, refer to ‘**GEMM WT epithelial annotation’**). To constrict a Markov graph, we considered the combined dataset of both WT and mutant–*Gfp* cells (refer to “**Pooled dataset of GEMM WT epithelial and GFP+ populations”)**. To ensure that our classification was specific to cell type related features, we performed feature selection, restricting genes to WT DEGs and excluding genes related to cell cycle, hypoxia, and apoptosis (*N*=234 genes). We projected sctranform residuals, restricted to these genes, onto the first 24 PCs selected by knee–point detection, corresponding to 90% variance. We obtained the first 8 diffusion components (DCs) retained by eigengap and transformed this diffusion graph into a Jaccard graph between *k*–neighborhoods using the Phenograph package (*21*). The resulting graph represented a Markov chain where we could calculate the Markov absorption probabilities for each unlabeled cell to reach a labeled cell of a given subtype. We initially considered six probabilities of basal (B) or each luminal (L1A, L1D, L1V, L1L, and L2). Given that we observed that mutant cell types largely demonstrated L2 or B probabilities, we summed the probabilities of all the L1 subsets (L1A, L1D, L1V, L1L) resulting in the sum of the L1 aggregated, L2, and B probabilities equaling to 1. We then assigned hard classification labels for the adenocarcinoma–*Gfp* cells by maximum likelihood of these subsets, findings supported by expression of canonical L1, L2, and B gene markers (**Figs. S7E)** and the set of published DEGs from L1, L2 and B signatures (from the AP lobe, **Figs S7D)** (*2*). For each set of DEGs, scanpy (*52*) score_genes function was used, which is the average z–scored expression of a set of B/L1/L2 genes (refer to **‘Human Genes Sets**) subtracted from the average z–scored expression of a randomly selected reference set of genes from a gene pool matched for binned expression values. We also considered the L1, L2, B to represent a deconvolution of mixed phenotypes that we represented by a 3–coordinate ternary graph (ggtern R package, Figs. 3A **and S7F**). As expected, the wild–type cells largely favored a single assignment of L1, B, and L2 even given mixed breeding backgrounds of WT mice. However, in both PtR and PtRP, adenocarcinoma cell probabilities became less committed to either B or L2, with several cells showing mixed B and L2 probabilities (Fig. 3A). Finally, to assess plasticity, we used the entropy of subtype probabilities per cell as a measure of basal–luminal mixing comparing wild–type to epithelial to mutant (PtR and PtPR) cells (statistic: Wilcoxon signed ranked test, right bar plot of Fig 3A). Of note, the majority of cells in NEPC, NEPC–P, *Tff3–Gfp* favored L2 probabilities with *Pou2f3–Gfp* and EMT–*Gfp* showing a mix L2 probabilities with either L1 or basal probabilities, respectively (**Fig. S7F**).

##### GEMM NEPC–A, NEPC–Ar and *Pou2f3–Gfp* classification

*Syp*+, *Chga*+, *Chgb*+, or *Pou2f3+* clusters (N=4,973 cells) were subsetted from all mutant epithelial cells (refer to ‘**GEMM Mutant*–Gfp* Cells PtR and PtRP annotation’)**. Normalized counts were projected onto the top 90 principal components (PCs) with 43% variance explained. Phenograph clusters (*k*=30) were identified yielding 17 clusters, which were merged based on the underlying hierarchical dendrogram. This yielded the following NEPC subsets *Pou2f3–Gfp* (3 clusters, 783 cells), *Ascl1* (NEPC–A, 12 clusters, 3516 cells), and *Ar* (NEPC–Ar, 2 clusters, 674 cells) (**Fig S9A and S9B**). We also noted three clusters which had shared expression of *Ar* and *Ascl1*, however these appeared hierarchically related to the *Ascl1* subset.

##### GEMM lymphoid and myeloid annotation

We subsetted CD45+ myeloid (monocyte, macrophages, PMNs, and DCs, 13,491 cells, *Cd14, Csfr1, Csfr3, Xcr1*). We projected sctransform residuals onto the top 84 principal components (PCs) detected by knee–point with 50% variance explained. We executed Phenograph clustering (*k*=30) yielding 25 clusters. We identified monocyte/macrophages (9,521 cells, 19 cluster, *Aif1, Csfr1*), polymorphonuclear neutrophils (3447 cells, 5 cluster, *Csfr3*), dendritic (523cells, 2 clusters, *Xcr1*) cells. We similarly subsetted CD45+ lymphoid (B cells, T cell, NK cells, 6,441 cells). We projected sctransform residuals onto the top 78 principal components (PCs) detected by knee–point with 35% variance explained. We executed Phenograph clustering (*k*=30), yielding 20 clusters. We identified NK/T cells (*Cd3e, Cd8, Cd4, Nkg7, Klrd1, Gzmb,* 19 clusters, 5,888 cells) and B cells (*Cd79a, Ms4a1,* 1 cluster, 553 cells) (Fig. 5B).

#### GEMM/Human Gene Sets

##### GEMM/Human gene sets

For AR scoring of cells, genes involved in androgen receptor signaling from published datasets (113 genes) (*53, 54*) were used. L1, L2 and basal cells were scored based on published DEGs. For each gene set, we computed a *per–cell*–score using the scanpy (*52*) score_genes function, which is the average z–scored expression of a set genes subtracted from the average z–scored expression of a randomly selected reference set of genes from a gene pool matched for binned expression values (*2*).

#### Identifying DEGs and Enriched Pathways

##### Identifying DEGs and MAST

For all differential expression, we used MAST *v.*1.16.0 (*55*). This package provides a flexible framework for fitting a hierarchical generalized linear model to the expression data. A regression model was executed that adjusted for both cellular detection rate (*cngeneson*, or number of genes detected per sample) and tissue status (Human: CSPC, CRPC, DNPC, NEPC) (GEMM: WT, PR 24, 30, 47 weeks, PRP 8, 9, 12, 16 weeks).

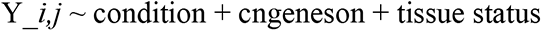

The condition represents the condition of interest and Y*i* is the expression level of gene *i* in cells of cluster *j*, transformed by natural logarithm with a pseudo–count of 1. We considered genes to be significantly differentially expressed for FDR <0.05 and absolute log fold–change (FC) > 0.3.

##### Human DEG comparisons

We performed differential expression (DEG) on the following: (1) Tumors cells labeled as CSPC, CRPC, DNPC, NEPC–A, and NEPC–N (refer to ‘**Human designation of adenocarcinoma and NEPC cells**’) *versus* rest, and NEPC–A *versus* NEPC–N (restricted to HMP17) (**Table S5**) and (2) each Phenograph cluster *versus* rest for each HP–CSPC and HMP–CRPC sample (select DEGs as dot plot shown in **Figs. S3–S4**).

##### GEMM DEG comparisons

We performed differential expression (DEG) on the following: (1) each GEMM cell types (B, L1, L2, Adeno–L1, Adeno–L2, Adeno–B, *Tff3*–*Gfp,* EMT–*Gfp, Pou2f3–Gfp*, NEPC *versus* rest **(Table S7)**, (2) each Phenograph cluster *versus* rest within the subset of wild-type and mutant cells (refer to ‘**Pooled dataset of GEMM WT epithelial and GFP+ populations’, Fig. S7B**) (3) and each GEMM NEPC subtype, NEPC–A, NEPC–Ar, and *Pou2f3–Gfp* **(Fig. S9D)**.

##### Identifying enriched pathways

Enriched gene pathways were identified using pre–ranked GSEA, as executed by the R package fGSEA (*56*) using 10,000 permutations. Gene ranks were calculated using –log(P–value) x log(FC) based on MAST differential expression (refer to **‘Identifying DEGs and MAST’**). To assess enriched pathways in between human cell types (CSPC, CRPC, DNPC, NEPC–A, and NEPC–N) or GEMM cell type, and each Phenograph cluster, we used a curated set of both prostate–specific (43 sets) (*2*) and developmental pathways (80 gene–sets) (*25, 26*) along with mouse cell type gene sets (67 gene sets) (*57*), KEGG (14 gene–sets), REACTOME (61 gene–sets), and HALLMARK (50 gene–sets) subset of canonical pathways in MSigBD*v*7.1 (*56*). All 315 included gene sets are listed in **Table S4**. We considered pathways with a P–value <0.05 to be significant. To score activation of significantly enriched pathways in our dataset, we would consider the leading–edge gene subset based on GSEA and calculated the average Z-score of the gene set.

#### Modeling Transitions Between Cell Types

##### GEMM pseudotime analysis

We aimed to model the transition of adenocarcinoma to its more transdifferentiated states (NEPC, *Pou2f3–Gfp*, EMT–*Gfp*, and *Tff3*–*Gfp* with a particular focus on the NEPC transition given its clinical relevance. We executed the Palantir (*30*) package to model these transdifferentiation processes. For the mapping of transcription factors across the adenocarcinoma and NEPC transition, we restricted to one model PtRP, which showed the majority of the NEPC cells. We ensured the results remained consistent across a number of robustness measures detailed in the **‘GEMM robustness of pseudotime’**. We add the caveat that the assumptions in Palantir may be problematic in cancer, specially that cells transition in one direction from the defined starting point and do no revert.

We first subsetted all mutant *Gfp*–positive cells (refer to **‘GEMM mutant*–gfp* cells PtR and PtRP annotation’,** N=21,499 cells), we projected normalized counts of the top 3000 highly variable genes (minus cell cycle) onto the first 95 PCs selected by knee–point explaining 48% of variance. Cells labeled as adenocarcinoma between PtR and PtRP showed good mixing, *albeit* most NEPC cells were contributed by the PtRP model (**Fig. S8A)**. Palantir uses diffusion maps constructed from a *knn* graph (*k*=30) to identify potential trajectories of transdifferentiation (*30*) thereby avoiding spurious edges due to the sparsity and noise of scRNA–seq. As input, we selected the first 14 diffusion components based on eigengap. We reasoned that a start cell from adenocarcinoma to more transdifferentiated states would display lower expression of low *Cdkn2a* expression (*Cdkn2a* upregulated as a result of *Trp53* and *Rb1* loss). This assumption was supported by the finding that NEPC and *Pou2f3–Gfp* cells showed the highest *Cdkn2a* expression, while wild type epithelial cells showed no *Cdkn2a* expression. To identify an appropriate *Cdkn2a*–low cell, we identified the Phenograph cluster of tumor cells with the lowest imputed median *Cdkn2a* (median imputed expression of 0.42 expression then randomly picked a cell within this cluster (start cell barcode: ‘PRP_9weeks_Intact_R1_157613955672309’). From this start cell, Palantir computed pseudotime and combined it with the neighbor–graph to construct a Markov chain that models state transitions as a stochastic process where an adenocarcinoma cell would reach more transdifferentiated states through a series of steps through the phenotypic manifold, whose probability is defined by the constructed Markov chain. We identified four ‘terminal’ states, *Pou2f3–Gfp*, NEPC, *Tff3–Gfp*, and EMT*–Gfp* **(Fig. S8A)**.

We then focused on the adenocarcinoma to NEPC transition given its clinical relevance. For the adenocarcinoma to NEPC transition, we subsetted 13,134 mutant *Gfp*–positive adenocarcinoma and NEPC cells restricting only to the PtRP model. We projected normalized counts of the top 3000 highly variable genes (minus cell cycle) onto the first 95 PCs selected by knee–point explaining 58% of variance. We again used the start cell barcode ‘PRP_9weeks_Intact_R1_157613955672309’’ (cluster 1) given the same reasoning as mentioned above and repeated pseudo–temporal ordering in its entirety. As an input, we selected the first 9 diffusion components based on eigengap. The terminal state was found within NEPC cluster 6 and centered around the cell ‘PRP_12weeks_Intact_R2_196638162925470.’ Gene trends for TF DEGs across the adenocarcinoma to NEPC transition were determined using a generalized additive model with cubic splines as the smoothing function across 500 equally sized bins, as executed by the Palantir package. Gene trends were grouped using Phenograph clusters (*k*=30) and we identified three TFs groups: early adenocarcinoma (purple), transition (red), and NEPC (blue) (Fig. 3D**, Figs. S8D)**. Heatmap of gene trends of select TF DEGs from each category are ordered by pseudotime modeling the transition from adenocarcinoma to NEPC colored by aforementioned groups (gene trends of imputed expression, scale –0.5 to 1.5) (Fig. 3E).

##### GEMM robustness of pseudotime

To ensure that our adenocarcinoma to NEPC pseudotime ordering was robust to cell sampling, we repeated our trajectory analysis by randomly subsampling cells 100 times. In particular, we subsampled 50% of cells from each Phenograph cluster (range 34 to 1647 cells) while randomly choosing a different cell each time in the low *Cdk2na* cluster (cluster 1, 1232 cells, **Fig. S8D, S8E**) We aimed to confirm that (1) the original pseudotime ordering of subsampled cells was highly correlated to newly generated pseudotime orderings based on Palantir analysis of the corresponding subsampled cells, (2) the original gene trends of DEGs of subsampled cells was highly correlated to newly generated gene trends of the corresponding subsampled cells, and (3) the terminal cell falls into the same end cluster with each run. Gene expression trends were computed using a generalized additive model of 8 splines with spline order 3 using the python package pyGAM (DOI: 10.5281/zenodo.1476122). Both pseudotime (**Fig. S8E in red**) ordering and gene trends were highly correlated between the original and subsampled runs (median Spearman correlation (*p*) for pseudotime ordering 0.98 and for all gene trends > 0.87). Moreover, the terminal cell fell in the original end cluster 6 in all 100 subsampled runs. Furthermore, putative transition genes (labeled in red, Fig. 3D) were highly expressed in Phenograph clusters that showed both mixed adenocarcinoma and NEPC states or high expression of *Cdk2na* in adenocarcinoma clusters **(Figs. S8F, S8G)**.

##### Human mapping genes across AR–high to AR–low–early–NEPC states using pseudotime ordering

To identify whether the set of transition TFs identified in the GEMM series showed similar trends in human CRPC samples, we reassessed our human scRNA–seq data to (1) establish the existence of an L2–like state and (2) understand the set of genes and pathways perturbed across this transition using the Palantir algorithm (*30*) **(Figs. 11C–E)**. We focused on CRPC–HMP13, a tumor that demonstrated areas of both low AR signaling and high AR signaling with gain of focal SYP–positive cells (Figs. 1D**, S3C**). We subsetted the tumor cells of HMP13 (N=354 cells), and projected normalized counts of the 5000 highly variable genes (removing cell cycle) onto the first 103 PCs selected by knee–point, which explained 52% of variance. Because pseudotime analysis can be sensitive to low cell sampling and may be problematic in cancer, we performed several robustness analyses as noted in section below **‘Human robustness of pseudotime.’**

Similar to the mouse pseudotime analysis (see “**GEMM pseudotime analysis”)** we used Palantir, and as input, we selected the first 16 diffusion components based on eigengap. We reasoned that a start cell of the trajectory from prostate adenocarcinoma to NEPC would display high AR signaling. To identify an appropriate AR–high start cell, we computed an AR–signaling–*per–cell*–score using the scanpy (*52*) score_genes function, which is the average z–scored expression of a set of AR target genes (refer to **‘GEMM/Human gene sets’**) subtracted from the average z–scored expression of a randomly selected reference set of genes from a gene pool matched for binned expression values. We then computed the mean AR score per Phenograph cluster, determining that cluster 0 (107 cells) had the highest AR score and randomly picked a start cell (start cell; HMP13_126777613212453). From this start cell, Palantir found a pseudo–temporal ordering from an AR–high cell to an AR–low–early–NEPC state through a series of steps through the phenotypic manifold. Only one branch probability was determined, in which the terminal state showed low PSA/FKBP5 state in the proximity of terminal cell: ‘HMP13_169150399240027’. Gene trends for DEGs across the AR–high to AR–low–early–NEPC transition were determined using a generalized additive model with cubic splines as the smoothing function across 500 equally sized bins, as executed by the Palantir package. Heatmap of gene trends of select transcription factor (TF) DEGs and mouse transition genes were ordered by pseudotime modeling the transition from AR–high to AR–low–early–NEPC transition (gene trends of imputed expression, scale –1 to 1.5) **(Fig S5C)**. Of note, cells with the highest L2 signature **(Fig. S11C)** had the high levels of JAK/STAT gene activation (cluster 1, blue) while those cells with early NEPC acquisition (cluster 2, green, refer to Fig. 3C for *SYP* expression) showed a relative downregulation in expression of JAK and STATs (**Fig. S11E**), consistent with our GEMM data.

##### Human robustness of pseudotime

To ensure that our adenocarcinoma to AR–high to AR–low–early–NEPC pseudotime ordering was robust to cell sampling, we repeated our trajectory analysis by randomly subsampling cells 100 times. In particular, we subsampled 50% of cells from each Phenograph cluster (range 97 to 150 cells) while randomly choosing a different cell each time in the high AR score cluster (cluster 0, 107 cells) We aimed to confirm that (1) the original pseudotime ordering of subsampled cells was highly correlated to newly generated pseudotime orderings based on Palantir analysis of the corresponding subsampled cells, (2) the original gene trends of DEGs of subsampled cells was highly correlated to newly generated gene trends of the corresponding subsampled cells, and (3) the terminal cell falls into the same end cluster with each run (**Fig. S11D**). Gene expression trends were computed using a generalized additive model of 8 splines with spline order 3 using the python package pyGAM (DOI: 10.5281/zenodo.1476122). Both pseudotime (**Fig. S11D in red**) ordering and gene trends were highly correlated between the original and subsampled runs (median Spearman correlation (*p*) for pseudotime ordering 0.84 and for all gene trends > 0.84 (except *SMARCC1* of 0.69, *ATF* 0.76, *STAT1* 0.79). Moreover, the terminal cell fell in the original end cluster 2 in all 100 subsampled runs.

#### Intersection of TFs Between GEMM and Human NEPC and SCLC

##### Shared phenotypes between GEMM NEPC, human NEPC, and human SCLC

We sought to assess the degree of shared phenotype 1) between mouse and human NEPC subtypes, and 2) between NEPC and SCLC. We therefore considered *ASCL1, NEUROD1,* and *POU2F3*–high subsets identified in our GEMM (NEPC–A and *Pou2f3–Gfp*) and human NEPC tumors (NEPC–A and NEPC–N), as well as corresponding SCLC subsets identified in a patient cohort of SCLC tumors (SCLC–A, SCLC–N, and SCLC–P) (*45*). For each NEPC subtype, we determined the set of significant DEGs using MAST with a FDR P–value ≤ 0.05 and coefficient ≥0.3 (≥0.2 in GEMM subsets) in a similar fashion to *Chan, et al.* (refer to Chan *et al.* Cancer Cell, 2021 ‘Differential Expression of Tumor and Immune Subsets’ SCLC–A, SCLC–N, and SCLC–P). From our GEMM dataset, we mapped significant DEGs to human orthologs. We then restricted our analysis to TFs based on the transcription factor database (TFDB, 16,65 total genes), leaving the following differentially expressed TFs with human orthologs for each subtype: GEMM NEPC (GEMM NEPC–A: 54 genes, GEMM *Pou2f3–Gfp*: 51 genes), human NEPC (Human NEPC–A: 36 genes, Human NEPC–N: 32 genes), and human SCLC (SCLC–A: 24 genes, SCLC –N: 22 genes, SCLC–P: 48 genes). We intersected the list of DEGs between subsets based on the following comparisons: 1) GEMM NEPC to human NEPC, 2) GEMM NEPC to human SCLC, and 3) human NEPC to human SCLC. The differences and intersections for human NEPC–A versus SCLC–A, human NEPC–N versus SCLC–N, GEMM NEPC–P versus SCLC–P were shown as Venn diagrams (Fig. 4E). A Fisher’s exact test was used to determine significance of overlap, considering only TFs from the TFDB.

To avoid hard thresholding of DEGs (MAST coefficient ≥ 0.3, see above, refer to ‘**Identifying DEGs and MAST’**) and further understand the relationship between neuroendocrine subtypes across lung and prostate cancer, we computed the *per*–gene (restricted to TFs) log_2_ fold change (FC) of mean Wasserstein distances (based on imputed expression, MAGIC, k=30, t=3) between *ASCL1 versus NEUROD1* or *POU2F3 versus* non-POU2F3 subtypes (**Fig. S7E**, left and right scatter plots, respectively). Specifically, for the *ASCL1* versus *NEUROD1* subsets, we first restricted our analysis to cells that were labeled as these subtypes (SCLC subtype labelling based on Chan *et al.* Cancer Cell, 2021) and to genes that are transcription factors (TF, TFDB, 16,65 total genes). For each TF gene, we calculated the Wasserstein distance between the *ASCL1* and *NEUROD1* labeled cell and calculated the log_2_FC of the mean Wasserstein distance (*ASCL1*/*NEUROD1)*. We then plotted the fold change for each cancer type, NEPC or SCLC, on the x– and y– axis, respectively. We further labeled TF genes that also overlapped DEGs (refer to ‘**Identifying DEGs and MAST’)** in blue and those with a FC ≥ 0.3 (enriched in *ASCL1*) and FC ≤ -0.3 (enriched in *NEUROD1*) both in NEPC and SCLC tumor types. We repeated these computational steps and calculations for the *POU2F3* subtype, but compared *POU2F3* labelled cells to all non–*POU2F3* subtypes.

#### Cell Type Signatures in Bulk RNA–seq Datasets from SU2C and PDX

##### SU2C and PDX bulk RNA–seq

To assess whether NEPC subtypes identified in the GEMMs are also present in patients in general, we considered publicly available bulk RNA–seq datasets from Stand Up to Cancer Prostate Cancer Foundation (SU2C–PCF) (*11*). Processed bulk RNA–seq expression data from 266 CRPC samples was downloade d from SU2C–PCF as Z–scored FPKM units per gene (https://github.com/cBioPortal/datahub/tree/master/public/prad_su2c_2019, file name: data_mRNA_seq_fpkm_polya_Zscores.txt,). Clinical annotations were also downloaded. Expression data was first queried for cell type markers (*ASCL1, POU2F3, NEUROD1, TFF3* and *ZEB2*) and corresponding signatures (NEPC–A, *Pou2f3–Gfp*, NEPC–N, *Tff3–Gfp* and EMT*–Gfp*). Corresponding signatures for the NEPC cell markers, NEPC–A (12 genes), *Pou2f3–Gfp* (10 genes), and NEPC–N (6 genes) were derived by taking the overlap of significantly expressed TF DEGs between NEPC and SCLC (refer to ‘**Shared phenotypes between GEMM NEPC, human NEPC, and human SCLC’).** For non–NEPC cell types, the top 50 DEGs for *Tff3–Gfp* and EMT*–Gfp* were used as the respective signatures (refer to ‘**Identifying DEGs and Enriched Pathway’**). A scatter plot was generated showing *Z*–scored gene expression in the SU2C database of *Pou2f3–Gfp* (y–axis) and NEPC–N (x–axis) signatures (Fig. 4D**, top**), and canonical transcriptional regulators *POU2F3* (y–axis) and *NEUROD1* (x–axis) (Fig. 4D**, bottom**). We also denoted non–NEPC or NEPC histology as annotated in the SU2C database by red and blue dots and Z–scored NEPC–A signature or ASCL1 expression by dot size. Similar plots were generated for *ZEB1* (marker of EMT, y-axis) and EMT signature (x-axis), or *TFF3* (y-axis) and TFF3 signature (x-axis) with dot size corresponding to Z–scored *ASCL1* expression **(Fig. S11B)**. Additionally, Dr. Peter Nelson provided log2 mean–centered ratios (Z–scores) of recently published and processed bulk RNA–sequencing data from 115 of PDX tumor samples (*32*). We specifically focused on *POU2F3* expression to identify PDX lines that may express *POU2F3*. We plotted the Z–scored *POU2F3* expression (x–axis) and *CHGA* (y–axis) expression with dot size corresponding to Z–scored *INSM1* expression. This analysis indicated two sublines derived from the same patient, one with NEPC–A (173_1_s6) and the other with *Pou2f3–Gfp* (173_2A_s11, 173_2B_s9, 173_2B_s11) phenotypes from two sublines derived from the same patient, LuCAP173. Notably, subline LuCAP173_2 (DNPC with rare foci of NEPC) showed high *POU2F3* expression, whereas subline LuCAP173.1 (fully NEPC) expressed low *POU2F3*, but high *INSM1, CHGA,* and *SYP* (**Fig. S11A**).

