## Supplementary Figure S1-S11 for "Multilineage plasticity in prostate cancer through expansion of stem–like luminal epithelial cells with elevated inflammatory signaling"

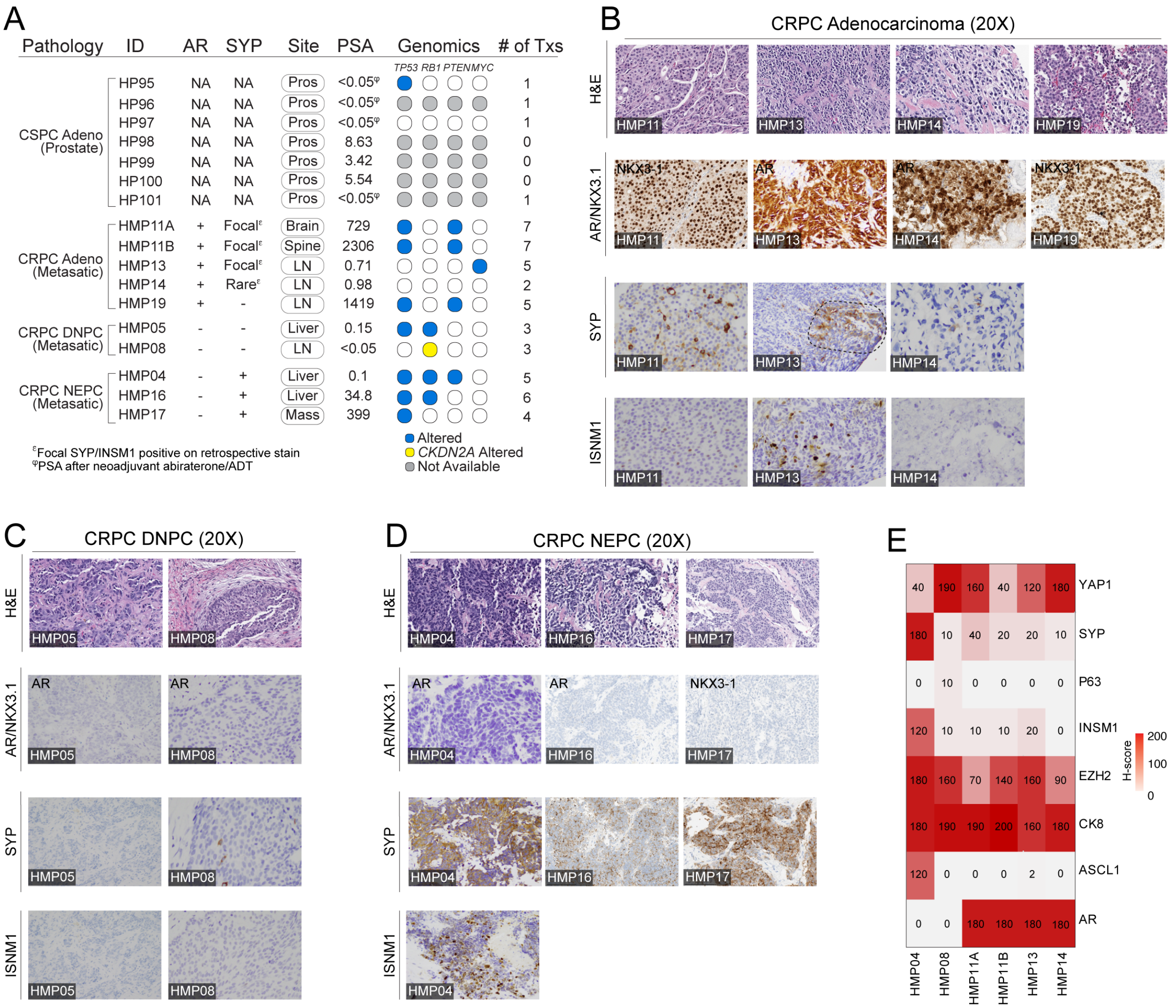

A
Pathology ID AR SYP Site PSA Genomics # of Txs
TP53 RB1 PTEN MYC
CSPC Adeno (Prostate)
HP95 NA NA Pros <0.05φ 1
HP96 NA NA Pros <0.05φ 1
HP97 NA NA Pros <0.05φ 1
HP98 NA NA Pros 8.63 0
HP99 NA NA Pros 3.42 0
HP100 NA NA Pros 5.54 0
HP101 NA NA Pros <0.05φ 1
CRPC Adeno (Metasatic)
HMP11A + Focalε Brain 729 7
HMP11B + Focalε Spine 2306 7
HMP13 + Focalε LN 0.71 5
HMP14 + Rareε LN 0.98 2
HMP19 + - LN 1419 5
CRPC DNPC (Metasatic)
HMP05 - - Liver 0.15 3
HMP08 - - LN <0.05 3
CRPC NEPC (Metasatic)
HMP04 - + Liver 0.1 5
HMP16 - + Liver 34.8 6
HMP17 - + Mass 399 4
εFocal SYP/INSM1 positive on retrospective stain
φPSA after neoadjuvant abiraterone/ADT
Altered
CKDN2A Altered
Not Available
B
CRPC Adenocarcinoma (20X)
H&E
HMP11 HMP13 HMP14 HMP19
AR/NKX3.1
NKX3-1 AR AR NKX3-1
HMP11 HMP13 HMP14 HMP19
SYP
HMP11 HMP13 HMP14
ISNM1
HMP11 HMP13 HMP14
C
CRPC DNPC (20X)
H&E
HMP05 HMP08
AR/NKX3.1
AR AR
HMP05 HMP08
SYP
HMP05 HMP08
ISNM1
HMP05 HMP08
D
CRPC NEPC (20X)
H&E
HMP04 HMP16 HMP17
AR/NKX3.1
AR AR NKX3-1
HMP04 HMP16 HMP17
SYP
HMP04 HMP16 HMP17
ISNM1
HMP04
E
YAP1: 40 190 160 40 120 180
SYP: 180 10 40 20 20 10
P63: 0 10 0 0 0 0
INSM1: 120 10 10 10 20 0
EZH2: 180 160 70 140 160 90
CK8: 180 190 190 200 160 180
ASCL1: 120 0 0 0 2 0
AR: 0 0 180 180 180 180
HMP04 HMP08 HMP11A HMP11B HMP13 HMP14
H-score
200
100
0

**Figure S1. Clinical, molecular, and pathological features of castrate–sensitive (CSPC) and castrate–resistant (CRPC) prostate cancer biopsies. (A)** Table of pertinent clinical and molecular features, including pathology group, sample ID, IHC for AR and INSM1, site of biopsy, PSA at time of biopsy, alterations in *TP53, RB1, PTEN,* and *MYC*, and number of prior treatments (also refer to **Table S2**). For hematoxylin and eosin (H&E) of primary CSPC samples, please refer to *Karthaus et al,* Science, 2020. **(B–D)** H&E and immunohistochemistry stains of AR, NKX3–1 or PSA, SYP and INSM1 for each patient biopsy (20X magnification) grouped by **(B)** CRPC adenocarcinoma (AR+SYP–), **(C)** CRPC double–negative prostate cancer (DNPC) (AR–SYP–), **(D)** CRPC neuroendocrine prostate cancer (NEPC) (AR+SYP+). For HP95, HP96, HP97, and HP101, PSA was <0.05 after neoadjuvant treatment with abiraterone and ADT (denoted by φ). Of note, HMP11A/B, HMP13 and HMP14 were found to have focal SYP on retrospective IHC for SYP, (denoted by ε) but were prospectively and clinically classified as AR+ CRPC adenocarcinoma. Only one representative biopsy is shown for HMP11A and HMP11B, which were derived pre– and post–enzalutamide (re–trialed with minimal/short response) treatment from a single patient. **(E)** Heatmap of *H*–score (immunohistochemical score, scale 0 to 200, refer to **Methods** for scoring) for select metastatic biopsies in those biopsies with ample archival tissue available.

Figure S2

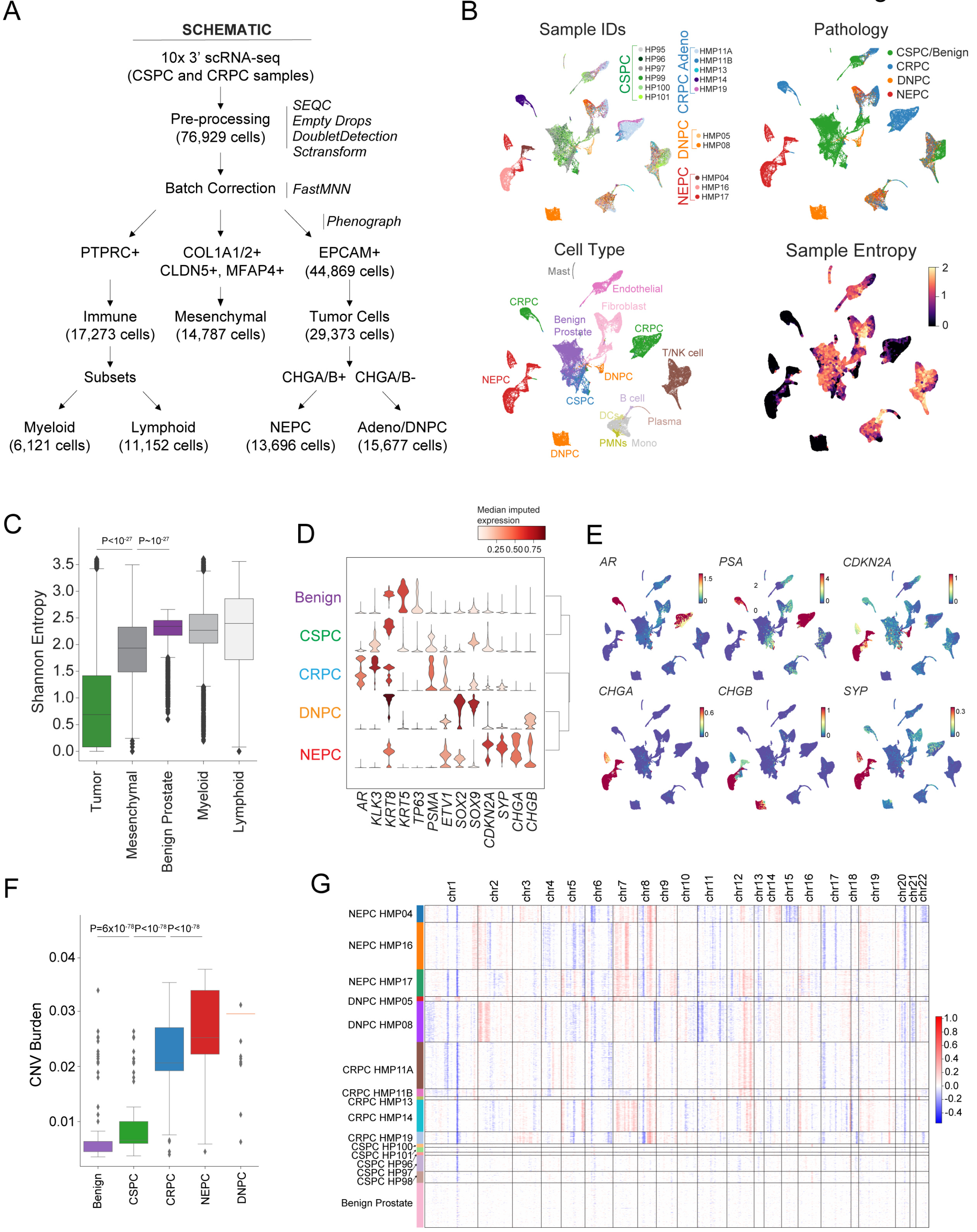
A
SCHEMATIC
10x 3' scRNA-seq
(CSPC and CRPC samples)
Pre-processing
(76,929 cells)
SEQC
Empty Drops
DoubletDetection
Sctransform
Batch Correction
FastMNN
Phenograph
PTPRC+
COL1A1/2+
CLDN5+, MFAP4+
EPCAM+
(44,869 cells)
Immune
(17,273 cells)
Mesenchymal
(14,787 cells)
Tumor Cells
(29,373 cells)
Subsets
CHGA/B+
CHGA/B-
Myeloid
(6,121 cells)
Lymphoid
(11,152 cells)
NEPC
(13,696 cells)
Adeno/DNPC
(15,677 cells)
B
Sample IDs
CSPC
HP95
HP96
HP97
HP99
HP100
HP101
CRPC Adeno
HMP11A
HMP11B
HMP13
HMP14
HMP19
DNPC
HMP05
HMP08
NEPC
HMP04
HMP16
HMP17
Pathology
CSPC/Benign
CRPC
DNPC
NEPC
Cell Type
Mast
Endothelial
Fibroblast
CRPC
Benign Prostate
T/NK cell
NEPC
DNPC
CSPC
B cell
DCs
Plasma
PMNs
Mono
Sample Entropy
C
P<10-27
P~10-27
Shannon Entropy
Tumor
Mesenchymal
Benign Prostate
Myeloid
Lymphoid
D
Median imputed expression
Benign
CSPC
CRPC
DNPC
NEPC
AR
KLK3
KRT8
KRT5
TP63
PSMA
ETV1
SOX2
SOX9
CDKN2A
SYP
CHGA
CHGB
E
AR
PSA
CDKN2A
CHGA
CHGB
SYP
F
P=6x10-78
P<10-78
P<10-78
CNV Burden
Benign
CSPC
CRPC
NEPC
DNPC
G
NEPC HMP04
NEPC HMP16
NEPC HMP17
DNPC HMP05
DNPC HMP08
CRPC HMP11A
CRPC HMP11B
CRPC HMP13
CRPC HMP14
CRPC HMP19
CSPC HP100
CSPC HP101
CSPC HP96
CSPC HP97
CSPC HP98
Benign Prostate
chr1
chr2
chr3
chr4
chr5
chr6
chr7
chr8
chr9
chr10
chr11
chr12
chr13
chr14
chr15
chr16
chr17
chr18
chr19
chr20
chr21
chr22

**Figure S2. Schematic, coarse cell typing, entropy and tumor annotation of human prostate cancer biopsies. (A)** Schematic for data processing, batch correction, cell type annotation and subsetting (refer to **Methods**). **(B)** UMAP plots (76,929 cells) of sample IDs grouped by CSPC (green shades, including benign prostate), CRPC adenocarcinoma (blue shades), DNPC (orange shades), and NEPC (red shades), pathology, cell type and sample entropy (refer to **Methods: 'Batch correction of human dataset'). (C)** Inter–patient heterogeneity measured by Shannon entropy based on tumor frequencies. To control for cell sampling, 100 cells were subsampled from each Phenograph cluster (*k*=30) within epithelial, immune, and mesenchymal compartments 100 times with replacement (Bonferroni–adjusted Student's t–test, **refer to Methods**). **(D)** Stacked violin plots of median imputed (MAGIC, *k*= 30, t=3) expression of genes associated with prostate cancer (*AR, KLK3/PSA, FOLH1/PSMA, ETV1, SOC2, SOX9, CDKN2A, SYP, CHGA, CHGB)* grouped by benign prostate, CSPC, CRPC, DNPC and NEPC, revealing higher expression of *PSA/KLK3*, *CDKN2A*, or *ETV1* in CSPC compared with benign cells. **(E)** Imputed (MAGIC, *k*=30, t=3) expression of *AR, KLK3/PSA, CHGA, CHGB, CDKN2A*, and *SYP* is shown on UMAP of all cells (N=76,929 cells). **(F)** Box plot of CNV burden per cluster (mean absolute value of CNV) stratified by benign and CSPC primary, CRPC, DNPC, and NEPC classified cells (Bonferroni–adjusted two sample t–test; refer to **Methods). (G)** Heatmap of *per* cell CNV estimated by sliding window approach (scale: modified expression –0.5 to 1, refer to **Methods**) with rows corresponding to cells grouped by sample and columns corresponding to ordered genomic location.

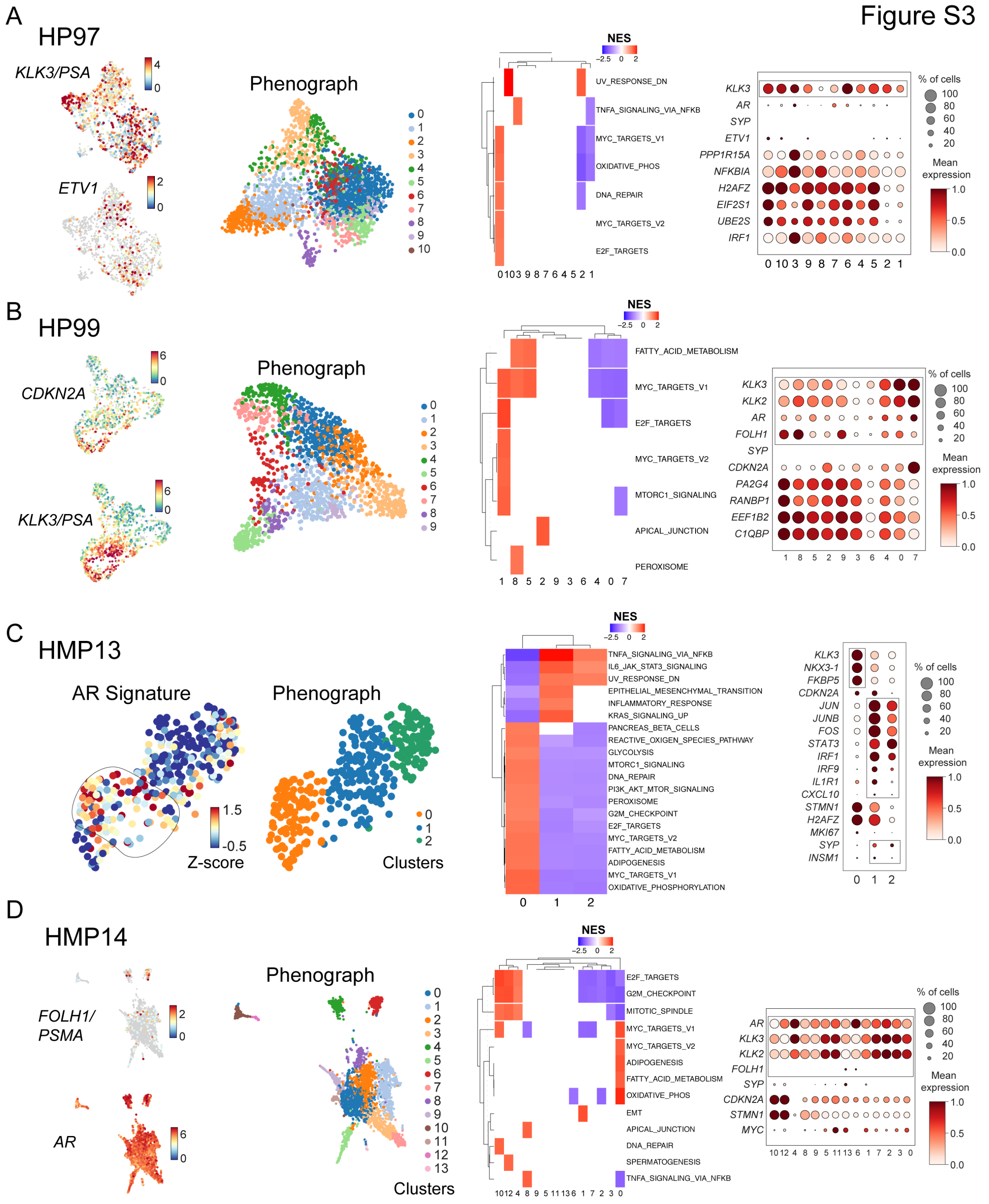

Figure S3
A
HP97
KLK3/PSA
ETV1
Phenograph
NES
UV_RESPONSE_DN
TNFA_SIGNALING_VIA_NFKB
MYC_TARGETS_V1
OXIDATIVE_PHOS
DNA_REPAIR
MYC_TARGETS_V2
E2F_TARGETS
KLK3
AR
SYP
ETV1
PPP1R15A
NFKBIA
H2AFZ
EIF2S1
UBE2S
IRF1
% of cells
Mean expression
B
HP99
CDKN2A
KLK3/PSA
Phenograph
NES
FATTY_ACID_METABOLISM
MYC_TARGETS_V1
E2F_TARGETS
MYC_TARGETS_V2
MTORC1_SIGNALING
APICAL_JUNCTION
PEROXISOME
KLK3
KLK2
AR
FOLH1
SYP
CDKN2A
PA2G4
RANBP1
EEF1B2
C1QBP
% of cells
Mean expression
C
HMP13
AR Signature
Z-score
Phenograph
Clusters
NES
TNFA_SIGNALING_VIA_NFKB
IL6_JAK_STAT3_SIGNALING
UV_RESPONSE_DN
EPITHELIAL_MESENCHYMAL_TRANSITION
INFLAMMATORY_RESPONSE
KRAS_SIGNALING_UP
PANCREAS_BETA_CELLS
REACTIVE_OXIGEN_SPECIES_PATHWAY
GLYCOLYSIS
MTORC1_SIGNALING
DNA_REPAIR
PI3K_AKT_MTOR_SIGNALING
PEROXISOME
G2M_CHECKPOINT
E2F_TARGETS
MYC_TARGETS_V2
FATTY_ACID_METABOLISM
ADIPOGENESIS
MYC_TARGETS_V1
OXIDATIVE_PHOSPHORYLATION
KLK3
NKX3-1
FKBP5
CDKN2A
JUN
JUNB
FOS
STAT3
IRF1
IRF9
IL1R1
CXCL10
STMN1
H2AFZ
MKI67
SYP
INSM1
% of cells
Mean expression
D
HMP14
FOLH1/PSMA
AR
Phenograph
Clusters
NES
E2F_TARGETS
G2M_CHECKPOINT
MITOTIC_SPINDLE
MYC_TARGETS_V1
MYC_TARGETS_V2
ADIPOGENESIS
FATTY_ACID_METABOLISM
OXIDATIVE_PHOS
EMT
APICAL_JUNCTION
DNA_REPAIR
SPERMATOGENESIS
TNFA_SIGNALING_VIA_NFKB
AR
KLK3
KLK2
FOLH1
SYP
CDKN2A
STMN1
MYC
% of cells
Mean expression

**Figure S3. Intra–tumoral transcriptomic programs in select CSPC and CRPC adenocarcinoma biopsies. (A–D)** UMAPs of individual CSPC (HP97, N=1,726 cells and HP99, N=1,296 cells) and CRPC (HMP13, N=354 cells and HMP14, 3,351 cells) biopsies. For each biopsy, cells are colored based on normalized expression ($log_2X+1$) for select genes or Phenograph clusters (*k=30)* (left). For HMP13 in panel C, an AR score was calculated using the scanpy score_genes function, which is the average Z–scored expression of a set of AR target genes (refer to **Methods: 'GEMM/Human Gene Sets'**) subtracted from the average Z–scored expression of a randomly selected reference set of genes from a gene pool matched for binned expression values (Z–score of –0.5 to 1.5). Shown per sample is a heatmap of significantly enriched gene sets *per* cell type (middle, scale from –2.5 to +2.5 of GSEA normalized enrichment score (NES), $P<0.05$, **refer to Methods**) and a dot plot of select differentially expressed genes (DEGs), stratified by Phenograph cluster (right, mean normalized $log_2X+1$ expression scaled from 0 to 1; dot size represents percent of cells expressing a given gene). Box outline highlights consistent or heterogeneous expression of tumor cell markers across Phenograph clusters. These are notable for HMP13 in panel C showing clusters with high levels of EMT markers, and HMP14 in panel D with *AR* positivity, but only focal *FOLH1/PSMA* expression.

Figure S4

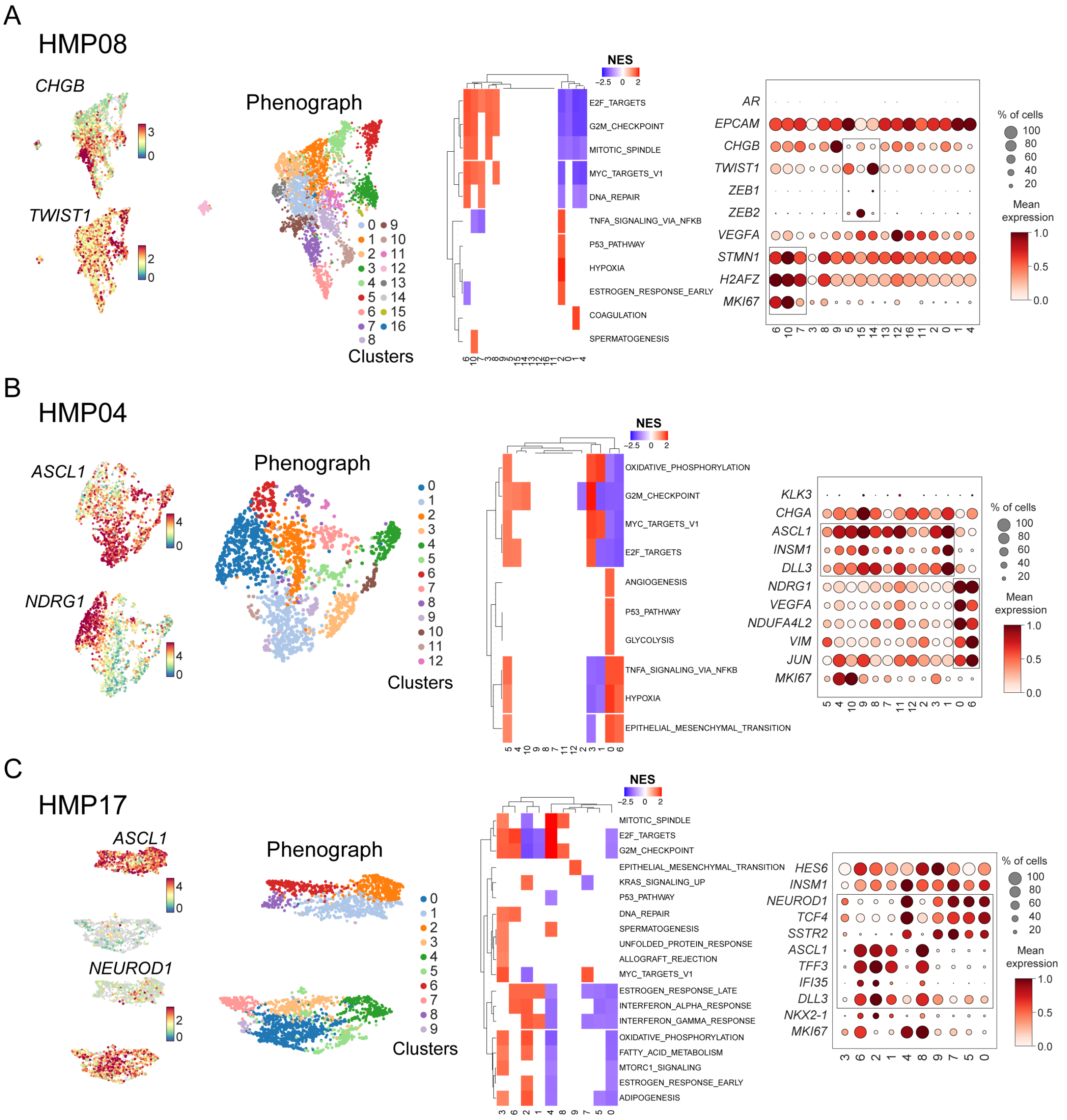
A
HMP08
CHGB
TWIST1
Phenograph
Clusters
NES
E2F_TARGETS
G2M_CHECKPOINT
MITOTIC_SPINDLE
MYC_TARGETS_V1
DNA_REPAIR
TNFA_SIGNALING_VIA_NFKB
P53_PATHWAY
HYPOXIA
ESTROGEN_RESPONSE_EARLY
COAGULATION
SPERMATOGENESIS
AR
EPCAM
CHGB
TWIST1
ZEB1
ZEB2
VEGFA
STMN1
H2AFZ
MKI67
% of cells
Mean expression
B
HMP04
ASCL1
NDRG1
Phenograph
Clusters
NES
OXIDATIVE_PHOSPHORYLATION
G2M_CHECKPOINT
MYC_TARGETS_V1
E2F_TARGETS
ANGIOGENESIS
P53_PATHWAY
GLYCOLYSIS
TNFA_SIGNALING_VIA_NFKB
HYPOXIA
EPITHELIAL_MESENCHYMAL_TRANSITION
KLK3
CHGA
ASCL1
INSM1
DLL3
NDRG1
VEGFA
NDUFA4L2
VIM
JUN
MKI67
% of cells
Mean expression
C
HMP17
ASCL1
NEUROD1
Phenograph
Clusters
NES
MITOTIC_SPINDLE
E2F_TARGETS
G2M_CHECKPOINT
EPITHELIAL_MESENCHYMAL_TRANSITION
KRAS_SIGNALING_UP
P53_PATHWAY
DNA_REPAIR
SPERMATOGENESIS
UNFOLDED_PROTEIN_RESPONSE
ALLOGRAFT_REJECTION
MYC_TARGETS_V1
ESTROGEN_RESPONSE_LATE
INTERFERON_ALPHA_RESPONSE
INTERFERON_GAMMA_RESPONSE
OXIDATIVE_PHOSPHORYLATION
FATTY_ACID_METABOLISM
MTORC1_SIGNALING
ESTROGEN_RESPONSE_EARLY
ADIPOGENESIS
HES6
INSM1
NEUROD1
TCF4
SSTR2
ASCL1
TFF3
IFI35
DLL3
NKX2-1
MKI67
% of cells
Mean expression

**Figure S4. Intra–tumoral transcriptomic programs in select DNPC and NEPC biopsies. (A–C)** UMAPs of individual DNPC (HMP08, N=4,279 cells) and NEPC (HMP04, N=1,791 cells and HMP17, N=2,845 cells) biopsies. For each biopsy, cells are colored based on normalized expression ($\log_2 X+1$) for select genes or number of Phenograph clusters (*k=30)* (left). Shown per sample is a heatmap of significantly enriched gene sets *per* cell type (middle, scale from –2.5 to +2.5 of GSEA NES, $P<0.05$, **refer to Methods**) and dot plot of select DEGs, stratified by Phenograph cluster (right, mean normalized $\log_2 X+1$ expression scaled from 0 to 1; dot size represents percent of cells expressing a given gene). Box outline highlights consistent or heterogeneous expression of genes of tumor cell markers across Phenograph clusters. These are notable for HMP08 in panel A showing high levels of EMT markers with focal *CHGB* expression, HMP04 and HMP17 in panels B and C showing *ASCL1* low, but *NRDG1* or *NEUROD1* high clusters, respectively.

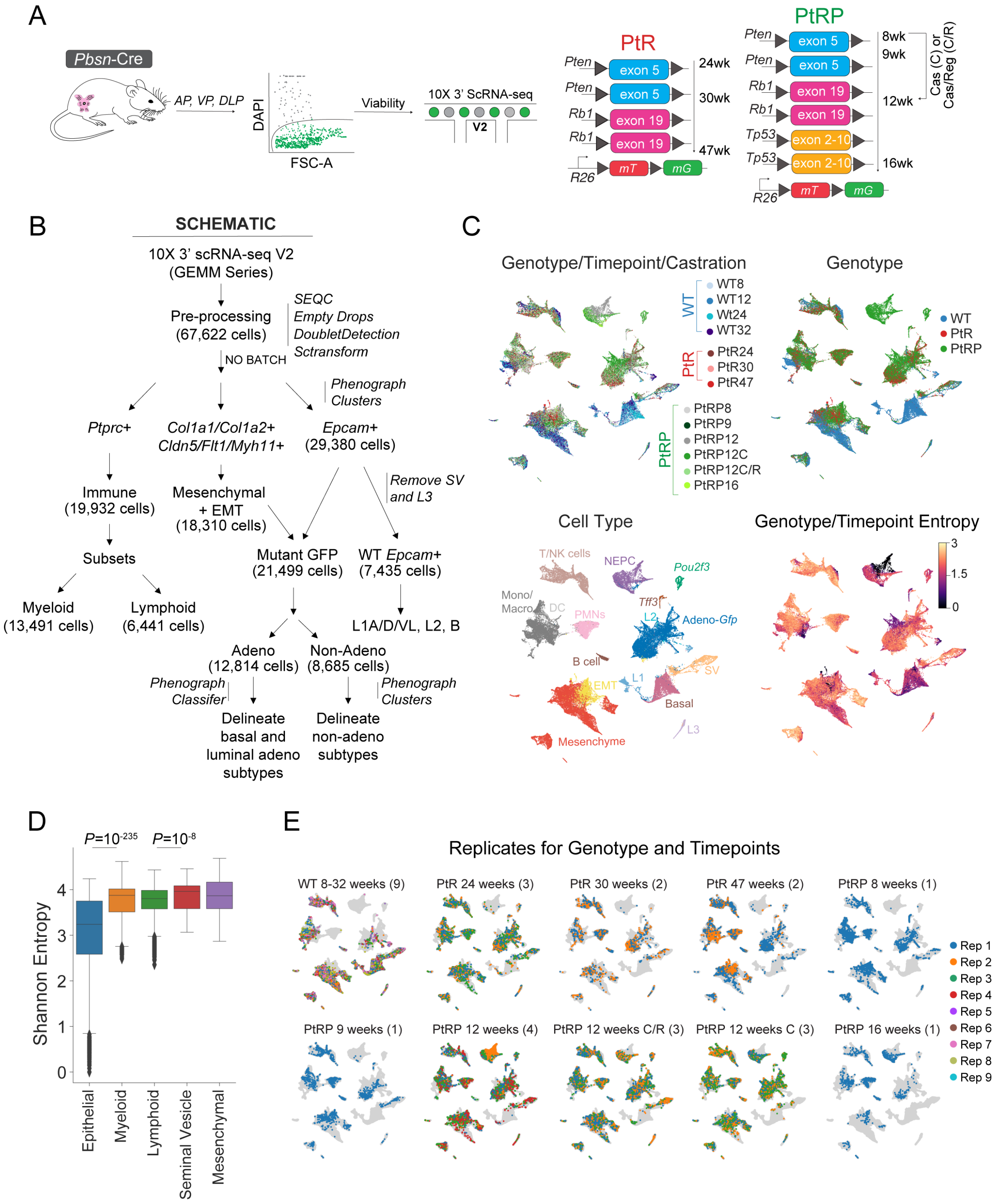
A
Pbsn-Cre
AP, VP, DLP
DAPI
FSC-A
Viability
10X 3' ScRNA-seq
V2
PtR
Pten exon 5
Pten exon 5
Rb1 exon 19
Rb1 exon 19
R26 mT mG
24wk
30wk
47wk
PtRP
Pten exon 5
Pten exon 5
Rb1 exon 19
Rb1 exon 19
Tp53 exon 2-10
Tp53 exon 2-10
R26 mT mG
8wk
9wk
12wk
16wk
Cas (C) or Cas/Reg (C/R)
B
SCHEMATIC
10X 3' scRNA-seq V2 (GEMM Series)
Pre-processing (67,622 cells)
SEQC
Empty Drops
DoubletDetection
Sctransform
NO BATCH
Phenograph Clusters
Ptprc+
Col1a1/Col1a2+ Cldn5/Flt1/Myh11+
Epcam+ (29,380 cells)
Immune (19,932 cells)
Mesenchymal + EMT (18,310 cells)
Remove SV and L3
Subsets
Mutant GFP (21,499 cells)
WT Epcam+ (7,435 cells)
Myeloid (13,491 cells)
Lymphoid (6,441 cells)
L1A/D/VL, L2, B
Adeno (12,814 cells)
Non-Adeno (8,685 cells)
Phenograph Classifer
Phenograph Clusters
Delineate basal and luminal adeno subtypes
Delineate non-adeno subtypes
C
Genotype/Timepoint/Castration
WT
WT8
WT12
Wt24
WT32
PtR
PtR24
PtR30
PtR47
PtRP
PtRP8
PtRP9
PtRP12
PtRP12C
PtRP12C/R
PtRP16
Genotype
WT
PtR
PtRP
Cell Type
T/NK cells
NEPC
Pou2f3
Mono/ Macro
DC
PMNs
Tff3
L2
Adeno-Gfp
B cell
EMT
L1
SV
Basal
L3
Mesenchyme
Genotype/Timepoint Entropy
3
1.5
0
D
P=10^-235
P=10^-8
Shannon Entropy
0
1
2
3
4
Epithelial
Myeloid
Lymphoid
Seminal Vesicle
Mesenchymal
E
Replicates for Genotype and Timepoints
WT 8-32 weeks (9)
PtR 24 weeks (3)
PtR 30 weeks (2)
PtR 47 weeks (2)
PtRP 8 weeks (1)
PtRP 9 weeks (1)
PtRP 12 weeks (4)
PtRP 12 weeks C/R (3)
PtRP 12 weeks C (3)
PtRP 16 weeks (1)
Rep 1
Rep 2
Rep 3
Rep 4
Rep 5
Rep 6
Rep 7
Rep 8
Rep 9

**Figure S5. Schematic, coarse cell typing replicates, and entropy of genetically engineered mouse models (GEMMs). (A)** Experimental design includes WT (9 mice), PtR (7 mice, *PbCre*:*Rosa26*$^{mT/mG}$*Pten*$^{fl/fl}$*Rb1*$^{fl/fl}$), and PtRP (13 mice, *PbCre*: *Rosa26*$^{mT/mG}$*Pten*$^{fl/fl}$*Rb1*$^{fl/fl}$*Tp53*$^{fl/fl}$) at labelled time points. Floxed exons for each gene are shown. All lobes (AP, VP, DLP) were digested followed by viability sort prior to scRNA–seq (refer to **Methods** for dissociation protocols). **(B)** Schematic for data processing, batch correction, cell type annotation and subsetting (refer to **Methods**). **(C)** UMAP plots (67,622 cells) of GEMMs grouped by genotype, timepoint, castration status (WT, blue shades; PtR, red shades; PtRP, green shades), genotype, cell type, and sample entropy (refer to **Methods: 'Batch correction'**). **(D)** Inter–sample heterogeneity measured by Shannon entropy based on sample (replicates treated as each sample) frequencies. To control for cell sampling, 100 cells were subsampled from each Phenograph cluster (*k*=30) within epithelial, immune, and mesenchymal compartments 100 times with replacement (Bonferroni–adjusted Student's t–test, **refer to Methods**). Seminal vesicles include both seminal vesicles and L3 epididymal cells. **(E)** UMAP plots of all cells (*N*=67,622 cells) colored by individual replicates *per* genotype, timepoint, and replicate. Numbers in parentheses in titles denote the number of replicates in each group.

Figure S6

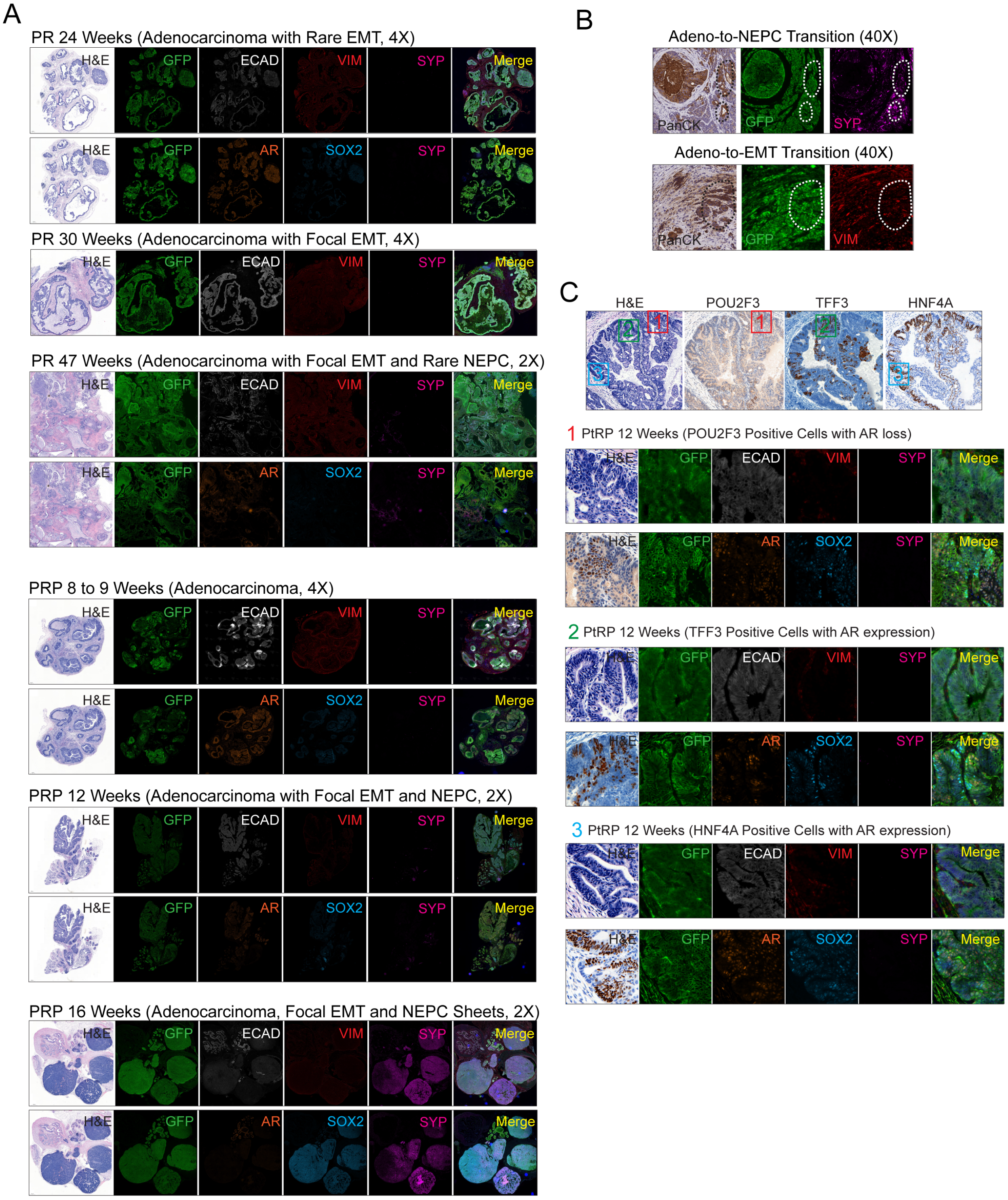
A
PR 24 Weeks (Adenocarcinoma with Rare EMT, 4X)
H&E
GFP
ECAD
VIM
SYP
Merge
AR
SOX2
PR 30 Weeks (Adenocarcinoma with Focal EMT, 4X)
PR 47 Weeks (Adenocarcinoma with Focal EMT and Rare NEPC, 2X)
PRP 8 to 9 Weeks (Adenocarcinoma, 4X)
PRP 12 Weeks (Adenocarcinoma with Focal EMT and NEPC, 2X)
PRP 16 Weeks (Adenocarcinoma, Focal EMT and NEPC Sheets, 2X)
B
Adeno-to-NEPC Transition (40X)
PanCK
GFP
SYP
Adeno-to-EMT Transition (40X)
VIM
C
H&E
POU2F3
TFF3
HNF4A
1 PtRP 12 Weeks (POU2F3 Positive Cells with AR loss)
2 PtRP 12 Weeks (TFF3 Positive Cells with AR expression)
3 PtRP 12 Weeks (HNF4A Positive Cells with AR expression)

**Figure S6. Spatial immunofluorescence in GEMM time course and confirmation of *Tff3*–, *Vim*–, *Pou2f3*–, and *Hnf4A*–positive cells. (A)** Validation of mutant subtypes for each genotype and time point by hematoxylin/eosin and multiplex immunofluorescence (mIF) (40x). Shown are GFP (mG, green), DAPI (nuclei, blue), VIM (mesenchymal, red), SYP (synaptophysin, purple), and E–CAD (e–cadherin, white). Magnification scales are noted (2X–4X). Majority of tumor type per genotype and time point is denoted above images. **(B)** Two regions of adeno–to–NEPC and adeno–to–EMT transition with PanCK IHC, and IF for GFP and SYP or VIM suggestive of transdifferentiation. **(C)** Hematoxylin/eosin and immunohistochemistry of POU2F3–, TFF3–, and HNFA– (4x) positive cells in mutant tissue adenocarcinoma. For spatial context, for POU2F3– (1, red label), TFF3– (2, green label), and HNFA– (3, blue label), hematoxylin/eosin, immunohistochemistry, and multiplex immunofluorescence (mIF) for GFP (mG, green), ECAD (white) or AR (orange), VIM (red) or SOX2 (turquoise), and SYP (synaptophysin, purple) are shown. Of note, while TFF3– and HNFA– co–stain with AR–positive cells, POU2F3–positive cells overlap with AR–negative cells.

Figure S7

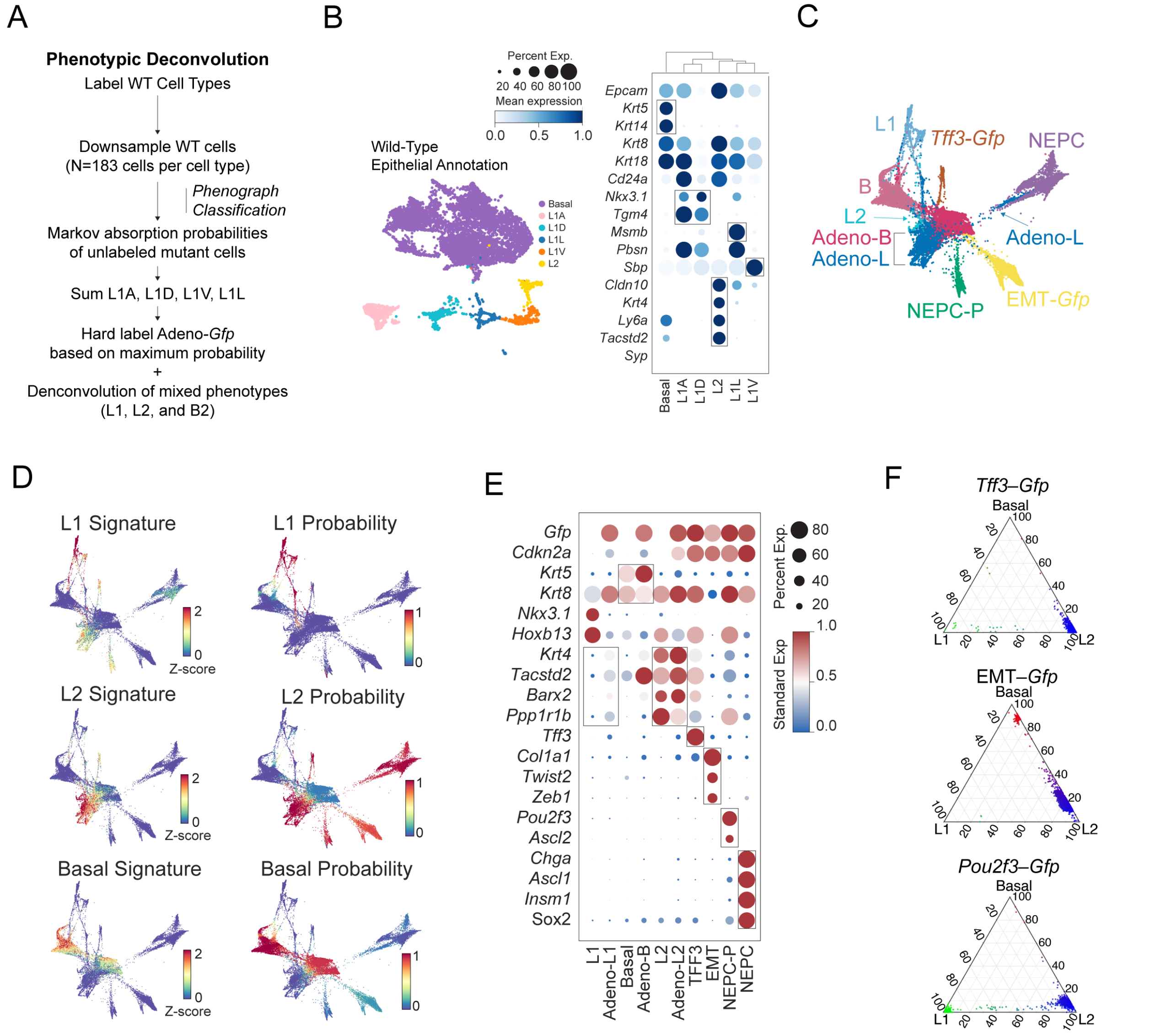
A
Phenotypic Deconvolution
Label WT Cell Types
Downsample WT cells
(N=183 cells per cell type)
Phenograph
Classification
Markov absorption probabilities
of unlabeled mutant cells
Sum L1A, L1D, L1V, L1L
Hard label Adeno-Gfp
based on maximum probability
+
Denconvolution of mixed phenotypes
(L1, L2, and B2)
B
Percent Exp.
20 40 60 80 100
Mean expression
0.0 0.5 1.0
Wild-Type
Epithelial Annotation
Basal
L1A
L1D
L1L
L1V
L2
Epcam
Krt5
Krt14
Krt8
Krt18
Cd24a
Nkx3.1
Tgm4
Msmb
Pbsn
Sbp
Cldn10
Krt4
Ly6a
Tacstd2
Syp
Basal
L1A
L1D
L2
L1L
L1V
C
L1
Tff3-Gfp
NEPC
B
L2
Adeno-B
Adeno-L
Adeno-L
NEPC-P
EMT-Gfp
D
L1 Signature
L1 Probability
L2 Signature
L2 Probability
Basal Signature
Basal Probability
Z-score
E
Gfp
Cdkn2a
Krt5
Krt8
Nkx3.1
Hoxb13
Krt4
Tacstd2
Barx2
Ppp1r1b
Tff3
Col1a1
Twist2
Zeb1
Pou2f3
Ascl2
Chga
Ascl1
Insm1
Sox2
L1
Adeno-L1
Basal
Adeno-B
L2
Adeno-L2
TFF3
EMT
NEPC-P
NEPC
Percent Exp.
80
60
40
20
Standard Exp
1.0
0.5
0.0
F
Tff3–Gfp
EMT–Gfp
Pou2f3–Gfp
Basal
L1
L2
20
40
60
80
100

**Figure S7. Phenotypic deconvolution of mutant cells in GEMMs. (A)** Schematic of phenotypic deconvolution of mutant *Gfp*–positive populations by calculating Markov absorption probabilities, as implemented in the Phenograph package. **(B)** UMAP of WT epithelial cells (N=7,345 cells, excluding seminal vesicles and L3 cells, **refer to Methods 'GEMM wild–type epithelial annotation')** with dot plot of select DEGs, stratified by cell type (mean normalized $\log_2 X+1$ expression scaled from 0 to 1; dot size represents percent of cells expressing the respective gene). **(C)** Force–directed layout (FDL) of wild–type and mutant epithelial cell types (N=28,934 cells) colored by wild–type and mutant cell types (N=28,934 cells) (also see **Fig. 2C**). **(D)** FDL scored using the scanpy score_genes function for published luminal 1 (L1), luminal 2 (L2) and basal (B) DEGs (left) or L1, L2, or B Markov absorption probabilities (right) (**refer to Methods: 'Phenotype deconvolution of GEMM *Gfp*–positive mutant cells')**. Note the correspondence for cell–specific signatures and absorption probabilities in Panel D with each wild–type and mutant cell type shown in Panel C. **(E)** Dot plot of select DEGs stratified by labelled and final cell type (mean normalized $\log_2 X+1$ expression scaled from 0 to 1; dot size represents percent of cells expressing a given gene). Outlining boxes highlight L2 markers (*Krt4, Tacstd2, Barx2, Ppp1r1b)*, or canonical basal *(Krt5)* or luminal *(Krt8)* markers. **(F)** Ternary plots of mutant *Tff3–Gfp,* EMT-*Gfp,* or *Pou2f3–Gfp* cells*,* based on three coordinates–basal (B), luminal 1 (L1), and luminal 2 (L2) Markov absorption probabilities. Each dot is colored by how likely the cell represents a B (red), L1 (green), or L2 (blue) phenotype. Corresponding plots of WT epithelial, mutant *Gfp*+, and NEPC cells are shown in **Fig. 3A**.

Figure S8

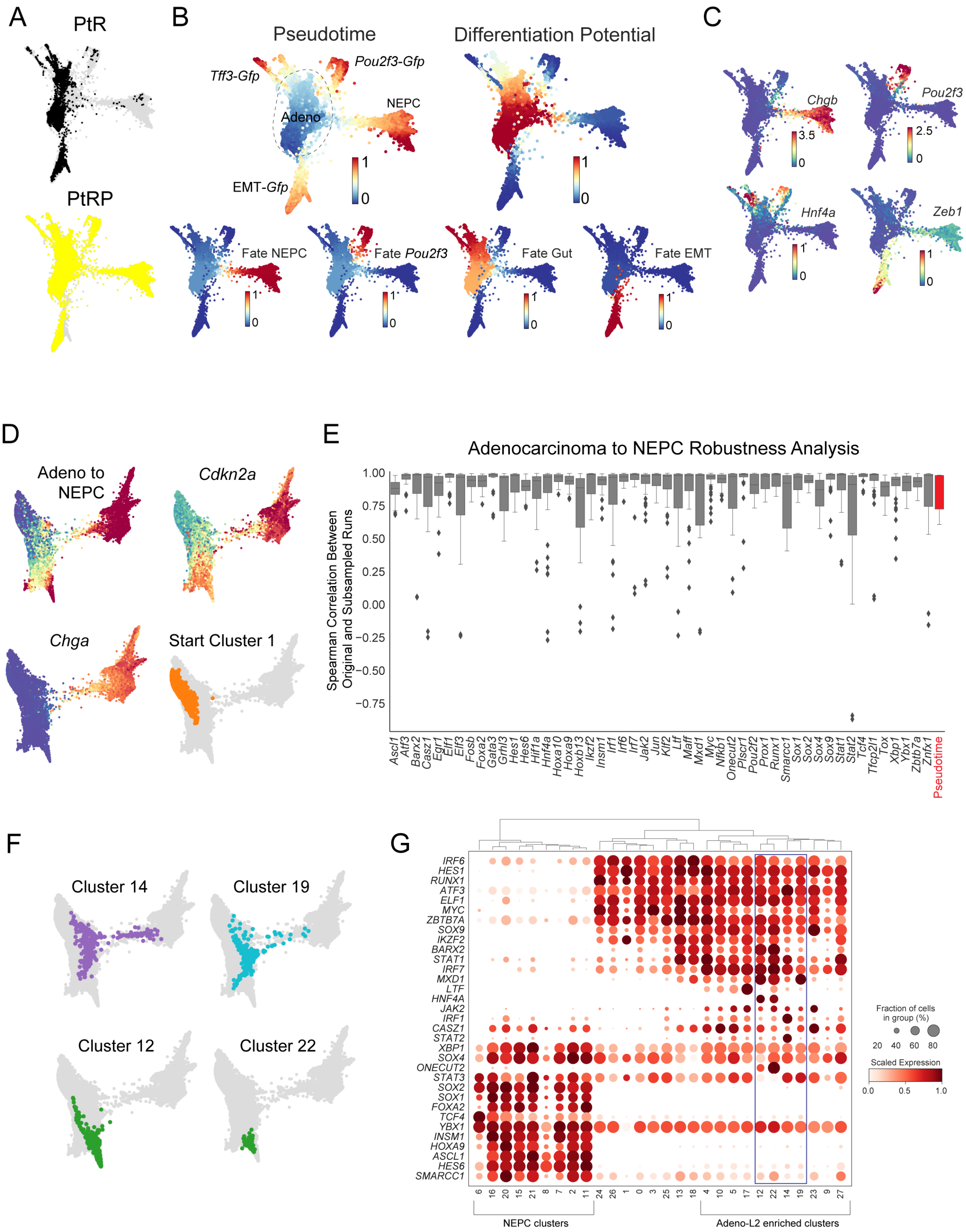
A
PtR
PtRP
B
Pseudotime
Differentiation Potential
Tff3-Gfp
Pou2f3-Gfp
Adeno
NEPC
EMT-Gfp
Fate NEPC
Fate Pou2f3
Fate Gut
Fate EMT
C
Chgb
Pou2f3
Hnf4a
Zeb1
D
Adeno to NEPC
Cdkn2a
Chga
Start Cluster 1
E
Adenocarcinoma to NEPC Robustness Analysis
Spearman Correlation Between Original and Subsampled Runs
Pseudotime
F
Cluster 14
Cluster 19
Cluster 12
Cluster 22
G
Fraction of cells in group (%)
Scaled Expression
NEPC clusters
Adeno-L2 enriched clusters

**Figure S8. Mapping transitions in adenocarcinoma to multi–lineage cell states in GEMMs. (A)** FDL of all mutant *Gfp*+ cells (refer to **Methods: 'GEMM Mutant–*Gfp* Cells PtR and PtRP annotation',** N=21,499 cells) colored genotype demonstrating overlap of adenocarcinoma cells between PtR and PtRP genotypes. **(B)** Shown is a FDL colored by pseudotime or differentiation potential (top) as implemented by Palantir with start cell randomly selected from the low *Cdkn2a* expression cluster (refer to **Methods: 'GEMM Pseudotime Analysis')**. 4 branch probabilities for either *Tff3–Gfp,* EMT–*Gfp, Pou2f3–Gfp*, or NEPC are also shown (bottom). **(C)** Imputed expression (MAGIC, k= 30, t=3) of *Chga, Zeb1, Pou2f3, and Hnf4a* are shown on FDL (N=21,499 cells, bottom). (**D)** To explore the clinically relevant adenocarcinoma to NEPC transition, mutant adenocarcinoma (subsetted to adeno–B and adeno–L2) and NEPCs cells, restricted to the PtR model, were subsetted (total N=13,134 cells**)**. FDL is colored by pseudotime scaled from 0 to 1, normalized $\log_2$X+1 expression of *Cdkn2a* or *Chga*, or start cluster (cluster 1 with lowest imputed median expression = 0.61) **(E)** Robustness of pseudotime analysis depicted by box plot of Spearman's rank correlations between observed and subsampled pseudotime results. Trajectory analysis was repeated 100 times while subsampling 50% of cells from each Phenograph cluster and randomly choosing a different start cell each time in the low *Cdk2na* cluster (cluster 1, 1232 cells). Red box (on right of box plot) denotes the distribution of Spearman's rank correlation between the observed and subsampled pseudo–temporal orderings. All other boxes denote the distribution of Spearman's rank correlation between the observed and subsampled gene trends. For each gene, expression trends were computed using a generalized additive model of 8 splines with spline order 3 using the python package pyGAM (refer to **Methods 'GEMM Robustness of Pseudotime').** Spearman's rank correlation was then calculated between the original and subsampled gene trends**. (F)** FDL of mutant adenocarcinoma (including adeno–B and adeno–L2) and NEPCs cells (total N=16,593 cells**)** showing four potential, and example, transition Phenograph clusters (*k*=30). Note clusters 14 and 19 contain both adenocarcinoma and NEPC cells. **(G)** Dot plot of select differentially expressed genes (DEGs) transcription factors, stratified by Phenograph cluster (right, mean normalized $\log_2$X+1 expression scaled from 0 to 1; dot size represents percent of cells expressing a given gene). Blue box outline highlights pertinent clusters shown in panel D. Clusters are also grouped by those enriched in Adeno–L2 and NEPC cells.

### Figure S9

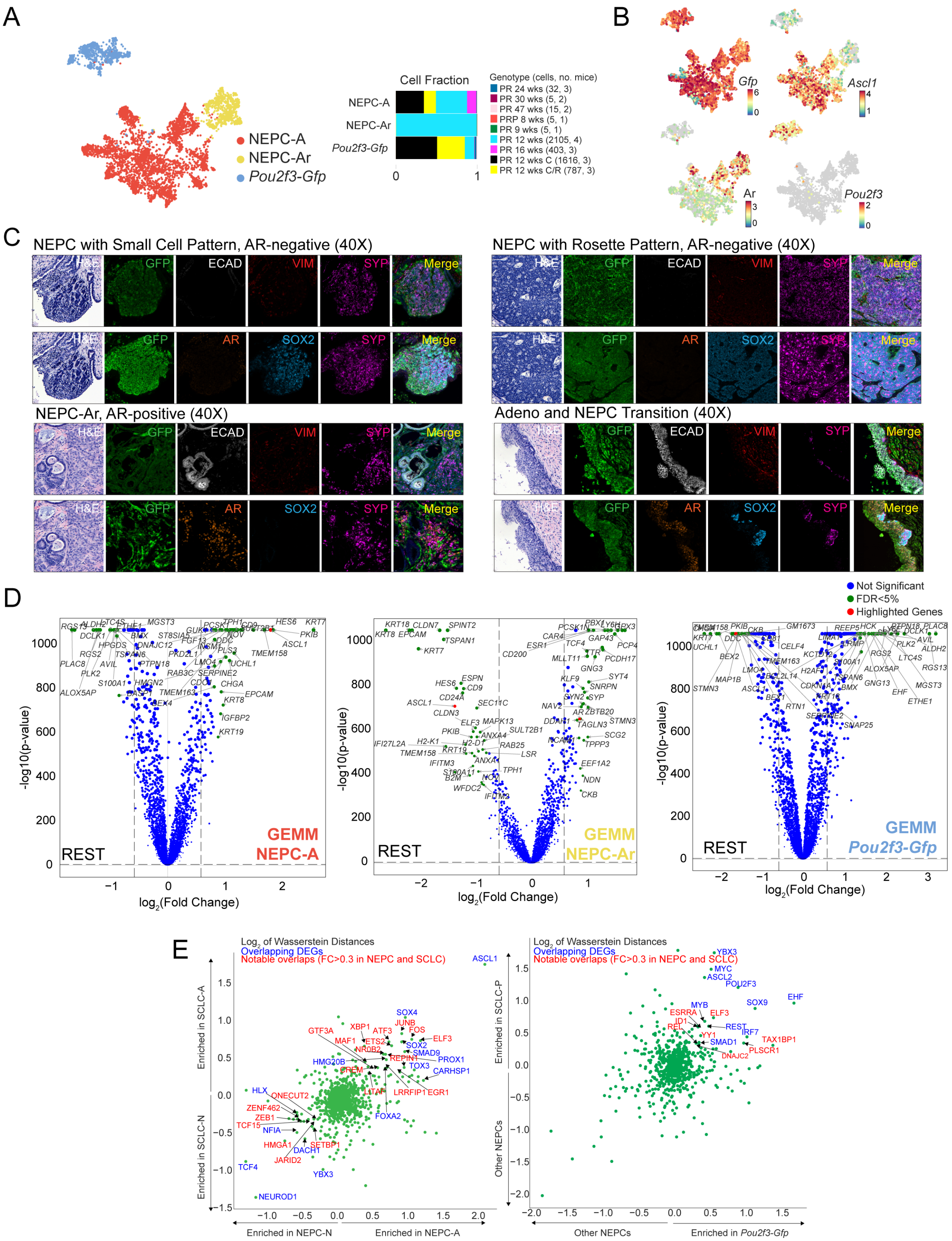

A
NEPC-A
NEPC-Ar
Pou2f3-Gfp
Cell Fraction
0
1
Genotype (cells, no. mice)
PR 24 wks (32, 3)
PR 30 wks (5, 2)
PR 47 wks (15, 2)
PRP 8 wks (5, 1)
PR 9 wks (5, 1)
PR 12 wks (2105, 4)
PR 16 wks (403, 3)
PR 12 wks C (1616, 3)
PR 12 wks C/R (787, 3)
B
Gfp
Ascl1
Ar
Pou2f3
C
NEPC with Small Cell Pattern, AR-negative (40X)
NEPC with Rosette Pattern, AR-negative (40X)
NEPC-Ar, AR-positive (40X)
Adeno and NEPC Transition (40X)
H&E
GFP
ECAD
VIM
SYP
Merge
AR
SOX2
D
Not Significant
FDR<5%
Highlighted Genes
-log10(p-value)
log2(Fold Change)
REST
GEMM NEPC-A
GEMM NEPC-Ar
GEMM Pou2f3-Gfp
E
Log2 of Wasserstein Distances
Overlapping DEGs
Notable overlaps (FC>0.3 in NEPC and SCLC)
Enriched in SCLC-A
Enriched in SCLC-N
Enriched in NEPC-N
Enriched in NEPC-A
Enriched in SCLC-P
Other NEPCs
Enriched in Pou2f3-Gfp

**Figure S9. Identification, characterization, and differential expression in NEPC subsets(A)** UMAP of GEMM mutant *Gfp*+ neuroendocrine cells colored by NEPC or *Pou2f3* subtypes (left; N=4,973 cells), based on expression of *Pou2f3* (*Pou2f3–Gfp*, 3 clusters, 783 cells), *Ascl1* (NEPC–A, 12 clusters, 3516 cells), and *Ar* (NEPC–Ar, 2 clusters, 674 cells). Subsets were determined by merging Phenograph clusters (*k*=30) based on the underlying hierarchical dendrogram (**refer to Methods**, shown on left). Bar plot shows fraction of cell types *per* genotype and time point for each NEPC or *Pou2f3–Gfp* subtype (shown on right). **(B)** UMAP plots colored by normalized expression $\log_2$X+1 of *Gfp, Ascl1, Ar, and Pou2f3.* Gray dots indicate zero expression. **(C)** Hematoxylin/eosin and multiplex immunofluorescence (mIF) (40x) characterizing NEPC–AR positive and negative subtypes, and transition populations. Shown are either GFP (mG, green), DAPI (nuclei, blue), VIM (mesenchymal, red), SYP (synaptophysin, purple), and E–CAD (e–cadherin, white); or GFP (mG, green), AR (androgen receptor, orange), SOX2 (transcription factor SOX2, cyan), and SYP (synaptophysin, purple). **(D)** Volcano plots of each NEPC or *Pou2f3* subtype *versus* rest of NEPC cells with genes represented by dots (y–axis, –log10 of MAST p–value FDR; x–axis $\log_2$ fold change). Green dots denote genes with FDR < 5% (refer to '**Identifying DEGs')**. Genes without significance are colored in blue. Select DEGs are noted in red dots. Dotted lines represent a fold change cut–off ≥ 1.5. **(E)** $\text{Log}_2$ fold change (FC) of mean Wasserstein distances, *per*–gene (restricted to transcription factors, TFs), comparing the imputed expression of NEPC–A *vs.* NEPC–A (shown on x–axis) or SCLC–A *vs.* SCLC–N (shown on y–axis). Genes with overlapping DEG TFs between either *ASCL1* or *NEUROD1* subset of SCLC and NEPC (refer to '**Shared phenotypes between GEMM NEPC, human NEPC, and human SCLC')** are shown in blue. TFs with $\log_2$ FC of greater ≥ 0.3 (enriched in *ASCL1*) or ≤ than -0.3 (enriched in *NEUROD1*) are shown in red (left). Similar scatter plot comparing *POU2F3* subset is shown, where the *POU2F3* subset is compared all non–*POU2F3* NEPC (x–axis) or SCLC (y–axis) subsets (right). These findings demonstrate conservation of transcription factors in NEPC subsets between prostate and lung cancer.

### Figure S10

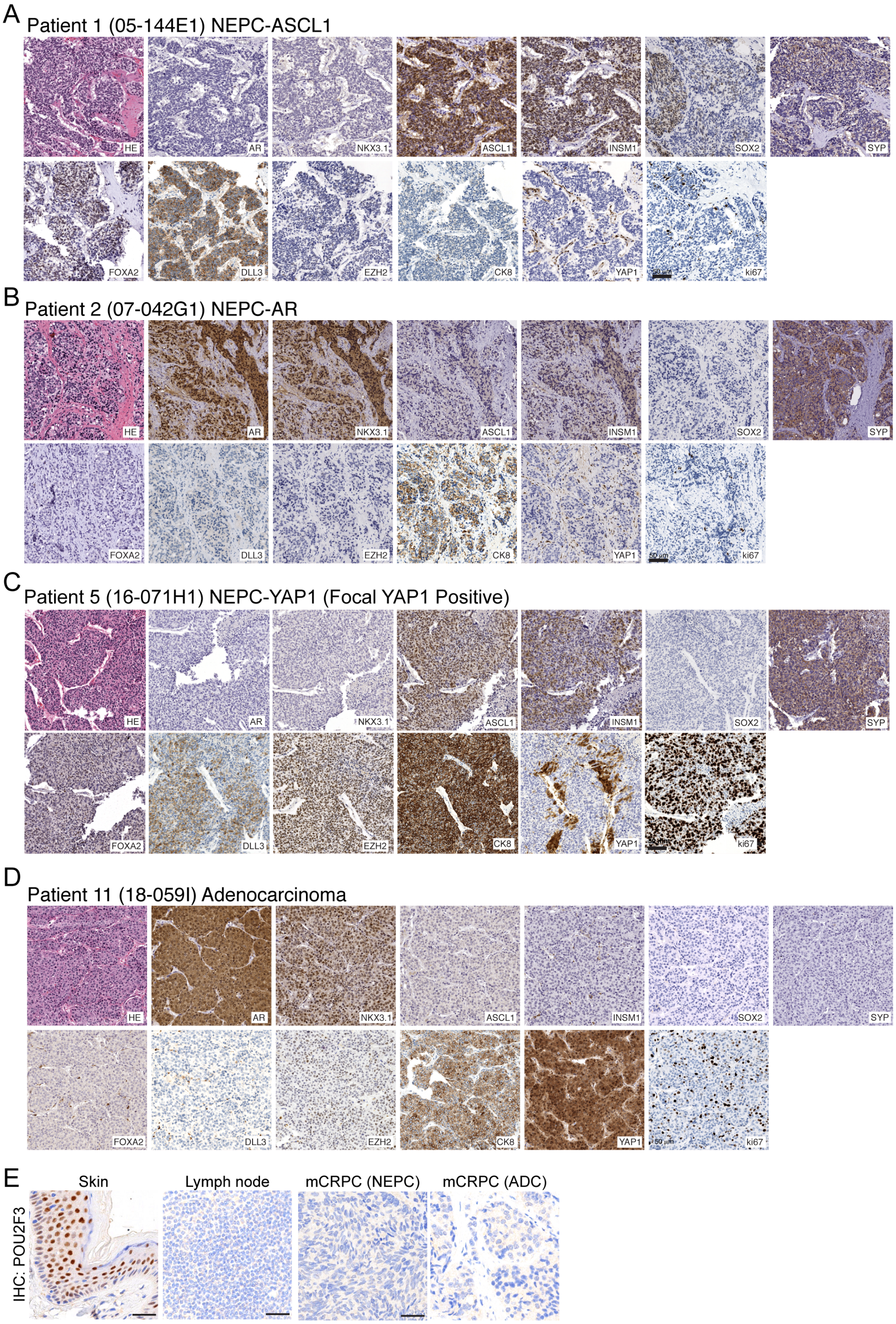
A
Patient 1 (05-144E1) NEPC-ASCL1
HE
AR
NKX3.1
ASCL1
INSM1
SOX2
SYP
FOXA2
DLL3
EZH2
CK8
YAP1
ki67
B
Patient 2 (07-042G1) NEPC-AR
HE
AR
NKX3.1
ASCL1
INSM1
SOX2
SYP
FOXA2
DLL3
EZH2
CK8
YAP1
ki67
C
Patient 5 (16-071H1) NEPC-YAP1 (Focal YAP1 Positive)
HE
AR
NKX3.1
ASCL1
INSM1
SOX2
SYP
FOXA2
DLL3
EZH2
CK8
YAP1
ki67
D
Patient 11 (18-059I) Adenocarcinoma
HE
AR
NKX3.1
ASCL1
INSM1
SOX2
SYP
FOXA2
DLL3
EZH2
CK8
YAP1
ki67
E
Skin
Lymph node
mCRPC (NEPC)
mCRPC (ADC)
IHC: POU2F3

**Figure S10. Extended select immunohistochemistry from human TMA data. (A)** Patient 1 (05–144E1) with neuroendocrine prostate cancer subtype ASCL1 (NEPC–A) with small cell morphology and high expression of SYP and ASCL1. INSM1 and DLL3 are also positive, while AR, CK8, NKX3.1, YAP1 and POU2F3 are negative. Expression of SOX2, FOXA2 and EZH2 are heterogeneous. **(B)** Patient 2 (07–042G1) with neuroendocrine prostate cancer amphicrine subtype (NEPC–AR) with poorly differentiated carcinoma morphology and expression of both SYP and AR. **(C)** Patient 5 (16–071H1) with neuroendocrine prostate cancer subtype Y (NEPC–Y) with poorly differentiated carcinoma morphology and expression of SYP and focal YAP1. Of note, most NEPC tumors showed YAP1 silencing with rare tumors demonstrating YAP1 focal and leaky expression. **(D)** Patient 11 (18–059I) with high grade prostate adenocarcinoma (ADC) with *NKX3.1* and YAP1, but lack of *SYP* or *INSM1* positivity. **(E)** POU2F3 immunohistochemistry with positive control (skin), negative control (lymph node), and no positivity in our CRPC NEPC or adenocarcinoma (ADC) TMA cohort.

Figure S11

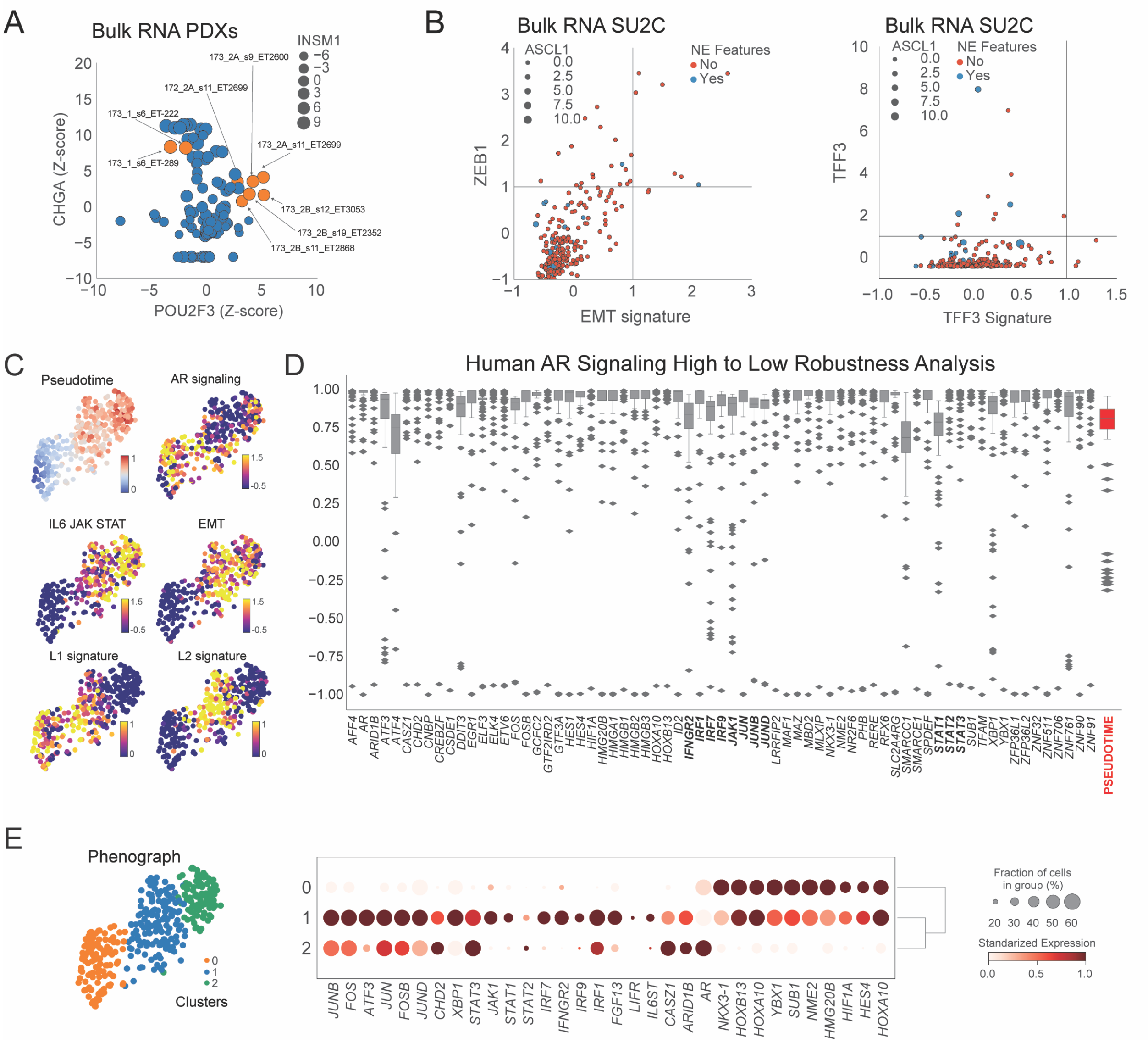

A
Bulk RNA PDXs
INSM1
CHGA (Z-score)
POU2F3 (Z-score)
B
Bulk RNA SU2C
ASCL1
NE Features
No
Yes
ZEB1
EMT signature
Bulk RNA SU2C
TFF3
TFF3 Signature
C
Pseudotime
AR signaling
IL6 JAK STAT
EMT
L1 signature
L2 signature
D
Human AR Signaling High to Low Robustness Analysis
PSEUDOTIME
E
Phenograph
Clusters
Fraction of cells in group (%)
Standarized Expression
JUNB
FOS
ATF3
JUN
FOSB
JUND
CHD2
XBP1
STAT3
JAK1
STAT1
STAT2
IRF7
IFNGR2
IRF9
IRF1
FGF13
LIFR
IL6ST
CASZ1
ARID1B
AR
NKX3-1
HOXB13
HOXA10
YBX1
SUB1
NME2
HMG20B
HIF1A
HES4
HOXA10

**Figure S11. Mapping of human TFs from AR–high to AR–low–early–NEPC states and evaluation of GEMM cell types in PDX and SU2C datasets.** **(A)** Scatter plot of Z–scored *POU2F3* (x–axis) and *CHGA* (y–axis) expression with dot size corresponding to Z–scored *INSM1* expression from bulk RNA–sequencing patient–derived xenograft (PDX) models of prostate cancer. This analysis indicated two sublines derived from the same patient, one with NEPC–A (173_1_s6) and the other with POU2F3 (173_2A_s11, 173_2B_s9, 173_2B_s11) phenotypes from two sublines derived from the same patient, LuCAP173 (highlighted in orange dots). **(B)** Scatter plots of gene signatures projected onto the SU2C bulk RNA–seq dataset. *Z*–scored EMT signature (top 50 DEGs) *versus* associated marker *ZEB1* (left). *Z*–scored TFF3 (50 DEGs) signature *versus* associated marker *TFF3* (right). Both EMT and TFF3 signatures were defined as the top 50 DEGs when comparing the given cell type to all other epithelial cell types using MAST (refer to **Methods: 'SU2C and PDX bulk RNA–seq'** and '**Identifying DEGs'**). Red and blue dots denote non–NEPC or NEPC features as annotated in the SU2C database, respectively. Dot size corresponds to the Z–score of ASCL1 expression. Dotted lines denote Z–score cutoff of 1 (refer to **Fig. 4D** for additional SU2C analyses). **(C)** UMAP of HMP13 (N=353 cells) colored by pseudotime or androgen receptor (AR), IL6–JAK–STAT, epithelial–to–mesenchymal transition (EMT), luminal 1 (L1) or luminal 2 (L2) signature scores. The AR, L1 or L2 signature scores were calculated using the scanpy score_genes function, which is the average Z–scored expression of a set of genes (refer to **Methods: 'GEMM/Human Gene Selection'** for AR target genes, L1 or L2 DEGs) subtracted from the average Z–scored expression of a randomly selected reference set of genes from a gene pool matched for binned expression values. For IL6–JAK–STAT or EMT scores, the Z–score of the leading edge of the respective HALLMARK GSEA term was used (refer to **Methods**). **(D)** Robustness of pseudotime analysis depicted by box plot of Spearman's rank correlations between observed and subsampled pseudotime results. Trajectory analysis was repeated 100 times while subsampling 50% of cells from each Phenograph cluster and randomly choosing a different start cell each time in the high AR score cluster (cluster 0, 107 cells, see panel C). Red box denotes the distribution of Spearman's rank correlation between the observed and subsampled pseudo–temporal orderings. All other boxes denote the distribution of Spearman's rank correlation between the observed and subsampled gene trends. For each gene, expression trends were computed using a generalized additive model of 8 splines with spline order 3 using the python package pyGAM (refer to **Methods** '**Human robustness of pseudotime').** Spearman's rank correlation was then calculated between the original and subsampled gene trends**. (E)** FDLs (N=353 cells) colored by 3 Phenograph clusters (*k*=30). (D) Dot plot of select transcription factors, stratified by Phenograph cluster (right, mean normalized $\log_2 X+1$ expression scaled from 0 to 1; dot size represents percent of cells expressing a given gene), highlighting activation JAK/STATs and interferon signaling genes along with genes involved in epithelial to mesenchymal transition.
